# The influence of visual cortex on perception is modulated by behavioural state

**DOI:** 10.1101/706010

**Authors:** Lloyd E. Russell, Zidan Yang, Pei Lynn Tan, Mehmet Fişek, Adam M. Packer, Henry W.P. Dalgleish, Selmaan Chettih, Christopher D. Harvey, Michael Häusser

## Abstract

Our understanding of the link between neural activity and perception remains incomplete. Microstimulation and optogenetic experiments have shown that manipulating cortical activity can influence sensory-guided behaviour or elicit artificial percepts. And yet, some perceptual tasks can still be solved when sensory cortex is silenced or removed, suggesting that cortical activity may not always be essential. Reconciling these findings, and providing a quantitative framework linking cortical activity and behaviour, requires knowledge of the identity of the cells being activated during the behaviour, the engagement of the local and downstream networks, and the cortical and behavioural state. Here, we performed two-photon population calcium imaging in L2/3 primary visual cortex (V1) of headfixed mice performing a visual detection task while simultaneously activating specific groups of neurons using targeted two-photon optogenetics during low contrast visual stimulation. Only activation of groups of cells with similar tuning to the relevant visual stimulus led to a measurable bias of detection behaviour. Targeted photostimulation revealed signatures of centre-surround, predominantly inhibitory and like-to-like connectivity motifs in the local network which shaped the visual stimulus representation and partially explained the change in stimulus detectability. Moreover, the behavioural effects depended on overall performance: when the task was challenging for the mouse, V1 activity was more closely linked to performance, and cortical stimulation boosted perception. In contrast, when the task was easy, V1 activity was less informative about performance and cortical stimulation suppressed stimulus detection. Altogether, we find that both the selective routing of information through functionally specific circuits, and the prevailing cortical state, make similarly large contributions to explaining the behavioural response to photostimulation. Our results thus help to reconcile contradictory findings about the involvement of primary sensory cortex in behavioural tasks, suggesting that the influence of cortical activity on behaviour is dynamically reassigned depending on the demands of the task.

---

Understanding the relationship between cortical activity and perception remains one of the most fundamental and challenging problems in neuroscience^1^. Microstimulation experiments have provided direct evidence for a causal role of activity in specific cortical circuits in biasing perception^2–7^. Moreover, microstimulation on its own can elicit artificial percepts^8–10^, as can optogenetic activation of cortical circuits^11–13^. Nevertheless, the number and functional identity of the stimulated neurons which are responsible for modulating behaviour are unknown^14,15^, although activity in just a single cell can be detected with extensive training^16–18^. Moreover, since the local and downstream network activity resulting from the manipulation have typically not been recorded, mechanistically linking the manipulation and behaviour via circuit dynamics has previously not been possible. Another complication is that silencing^19–21^ and lesion^22–25^ experiments have in some cases produced contradictory findings about the requirement for cortical activity in perception and behaviour^26,27^. The modulation of cortical responses by behavioural state^28–31^, task outcome^32^ and task demands^33,34^ have been well reported. However, how the modulation of cortical activity by state or task corresponds to the influence of that area on behaviour has only been studied using largely correlative methods^35,36^. Consequently, we lack a causal framework for linking activity in specific cortical populations with perception in different behavioural states.

To probe the importance of the identity of individual members in an active population of neurons and their influence on cortical activity and behaviour, we activated specific groups of cells distributed through a volume of visual cortex with two-photon optogenetic stimulation^37–39^ while performing simultaneous two-photon population calcium imaging of the same volume^40–45^. We employed our all-optical approach in mice trained on a visual detection task where task difficulty, and evoked cortical responses, were titrated by adjusting stimulus contrast. This allowed us to address the following questions. First, how important is the functional identity of individual members of an active group of cells? And second, under what conditions does cortical activity guide a perceptually-driven behaviour, and how does this depend on the specific routing of information through the local V1 circuit?

We coexpressed the calcium sensor GCaMP6s^46–48^ with the excitatory, somatically-restricted opsin C1V1 ^49,50^ in pyramidal cells of L2/3 primary visual cortex (V1) of mice performing a visually guided behaviour. Mice were head-fixed and trained to perform a visual stimulus detection task (**Fig. 1a**) incorporating a randomised nolick period before a small drifting sinusoidal grating patch with a random orientation was presented, during which a reward could be obtained via a lick spout (**Fig. 1b**). Mice learned the task quickly with a maximal contrast stimulus, reaching a high level of stable performance (**Supplementary Fig. 1**). Lowering the stimulus contrast modulated performance on the task (**Supplementary Fig. 1**). To test the influence of activity in functionally defined groups of neurons in V1 we targeted multiple cells for two-photon photostimulation while recording the resulting neuronal and behavioural responses. We first identified sensory- and photostimulation-responsive neurons in the retinotopically appropriate field of view (**Fig. 1d**), and then designed and stimulated three different ensembles of neurons (**Fig. 1e**): 1. Cotuned (CT, in which all constituent cells responded preferentially to the orientation of the low-contrast visual stimulus), 2. Non-cotuned (NCT, in which all constituent cells were responsive to the visual stimuli but preferred different orientations, with each of the 4 orientations represented equally) and 3. Non-responsive (NR, where all constituent cells were not responsive to the presented visual stimuli). We randomised the photostimulation of the three different ensemble types during low contrast visual stimulation with an orientation matched to the cotuned ensemble’s preference. We also presented the low-contrast visual stimulus without photostimulation and included trials where we stimulated the same ensembles in the absence of a visual stimulus. These trials were all interleaved with a high rate of non-photostimulation, high contrast, random-orientation trials to maintain engagement in the overall task. Averaged across all sessions in all mice, the detectability of the low contrast visual stimulus was unchanged by photostimulation (**Fig. 1f,g**, P(Lick) for Low: 0.35 ± 0.21, Low+CT: 0.34 ± 0.15, Low+NCT: 0.39 ± 0.19, Low+NR: 0.36 ± 0.19. *P* = 0.76 Kruskal-Wallis test. *N* = 21 sessions, 14 mice). Surprisingly, on a session by session basis, the stimulus detection rates on trials with photostimulation displayed a clear dependence on task performance during that session: mice performing poorly on low contrast trials were helped by photostimulation, and mice which were performing well were hindered by photostimulation (**Fig. 1h**, R^2^ = 0.48, *P* < 0.001 for CT stimulation). This relationship was observed when stimulating any of the three ensembles (**Supplementary Fig. 2**), but it reached statistical significance (in comparison to resampling the non-stimulation trials) only when stimulating the CT ensemble (**Fig. 1i**, slopes for CT: 0.52 ± 0.12, *P* = 0.004; NCT: 0.76 ± 0.12, *P* = 0.353; NR: 0.72 ± 0.13, *P* = 0.214; compared to slopes from the resampled distribution [0.86 ± 0.12]. Intercepts for CT: 0.16 ± 0.05, *P* = 0.010; NCT: 0.13 ± 0.05, *P* = 0.054; NR: 0.10 ± 0.05, *P* = 0.185; compared to intercepts from the resampled distribution [0.05 ± 0.05]. *N* = 21 sessions, 14 mice). We therefore focus the remainder of our analysis on the change in behaviour when stimulating CT ensembles.

**Figure 1.**
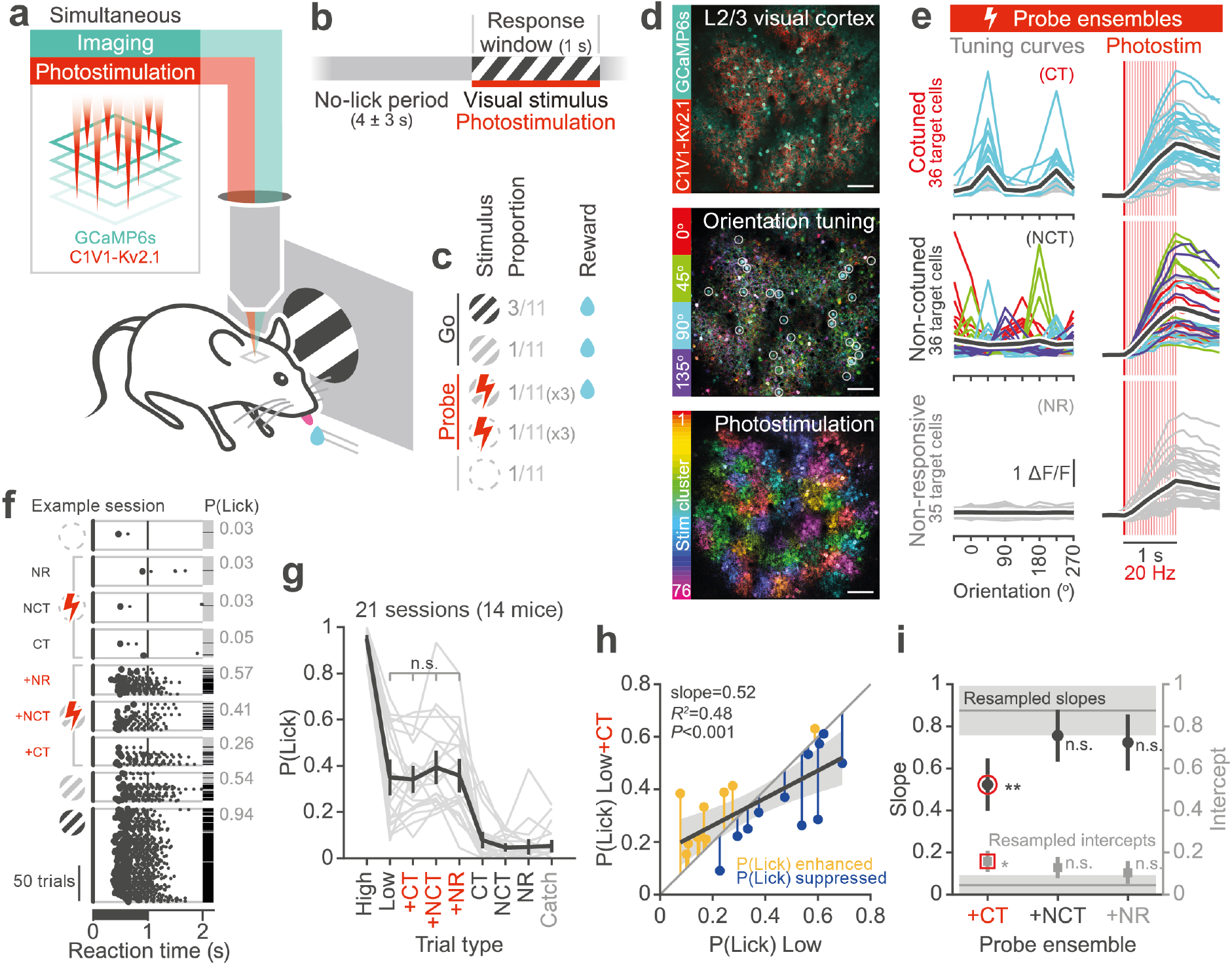
Two-photon photostimulation of cotuned ensembles influences detection behaviour depending on performance in the task.

**a.** Schematic outlining the experiment. Mice coexpress GCaMP6s and C1V1-Kv2.1 permitting cellular resolution reading and writing of neural activity. Mice are headfixed and are trained to perform a visual stimulus detection task. **b.** Structure of behavioural trials. After withholding licks for a randomised interval (4 ± 3 sec) a stimulus is presented to the mouse. The mouse can respond throughout the stimulus duration (1 sec) to receive a water reward. **c.** 5 different trial types are presented to the mouse in a pseudorandom blocked structure. High contrast trials are interleaved with low contrast, probe (photostimulation) and catch (no stimulus) trials. There are 3 types of stimulation ensemble for each probe trial type, giving a total of 9 different trial types per session. Any trial with a visual stimulus is rewarded if the mouse responds during the response window. **d.** Top: Example FOV (one plane from a 4-plane volume) showing construct expression in L2/3 mouse primary visual cortex. GCaMP6s is expressed transgenically and C1V1-Kv2.1 is expressed virally through injection. Middle: Visual stimulus orientation preference map. 4 different orientations of drifting gratings are presented to the mouse. Pixel intensity is dictated by the stimulus triggered average response magnitude. Hue corresponds to stimulus orientation. Bottom: Prior to designing the functionally defined stimulation ensembles we had to find which cells were expressing both constructs sufficiently for photostimulation. The majority of recorded cells were grouped into 76 different clusters of 50 cells each (distributed across 4 planes) and targeted for sequential photostimulation to confirm responsivity prior to the experiment. Pixel intensity indicates the change in florescence caused by photostimulation. Colour corresponds to the photostimulation cluster which caused the largest change in activity. White circles in the middle panel indicate example targets within this plane selected for targeted photostimulation of a cotuned ensemble. All scale bars 100 μm. **e.** 3 different types of stimulation ensemble are designed per experiment: cotuned (CT, all cells prefer same visual stimulus), non-cotuned (NCT, cells prefer different visual stimuli), and non-responsive (NR, cells are not responsive to the visual stimuli). Left: example visual stimulus orientation tuning curves of the target cells. Right: example average photostimulation response for all cells (coloured lines) and group average (black line) when only that group of cells is stimulated. Line colours indicate preferred orientation of visual stimulus for that cell. **f.** Example lick raster. Trials are sorted by type for display. The stimulus is delivered at time zero. Licks are indicated by black dots with the first lick indicated by a larger dot. Trial outcome is indicated to the right (Black = licked, grey = no lick). The average probability of licks per trial type for this example session is shown on the right. **g.** Average performance for all trial types across all animals (N = 14 mice, 21 sessions total). **h.** A strong relationship of behavioural modulation by CT photostimulation with task performance is seen. At low performances photostimulation enhances behavioural stimulus responses while at higher performances photostimulation suppresses responses. Diagonal unity line is shown. Grey shaded region indicates CI of fit. **i.** To account for possible regression to mean confounds the relationship in **h.** is compared to the range of expected fits from a resampling procedure using the mean lick probability and trial numbers for each set of low contrast trials. Only stimulation of cotuned ensembles results in a significant and detectable deviation from the permutation test bounds. Error bars indicate the standard error of the slope and intercept estimates.

The opposite effects of photostimulation depending on overall performance suggests that V1 serves different roles during different behavioural demands and cortical states. We therefore investigated the relationship between the state of cortex and the behavioural effects of cortical stimulation. We first examined network synchronicity, which is linked to arousal and attention^51,52^, in the period when the mice were waiting for a visual stimulus (**Fig. 2a**, measured by average pairwise correlations between all cells in the 4 seconds prior to the low contrast visual stimulus). We found that successful (hit) low-contrast visual stimulus trials were preceded by more asynchronous activity patterns than miss trials (**Fig. 2b**. Z-scored pre-trial correlation coefficient on hits: −0.35 ± 0.33, on misses: 0.05 ± 0.18, *P* = 0.0017 Wilcoxon sign rank test. *N* = 21 sessions, 14 mice) as recently reported ^53,54^. Since cortical synchronicity is linked to perceptual performance, we next asked if the average difference in pre-stimulus synchronicity between hit and miss trials in a given session varies with the level of overall performance in that session. We found evidence for a greater difference between the pre-stimulus network synchronicity on hit and miss low-contrast trials when the task was more difficult for the mice (**Fig. 2c**, *R^2^* = 0.18, *P* = 0.052. *N* = 21 sessions, 14 mice. See also **Supplementary Fig. 3**).

**Figure 2.**
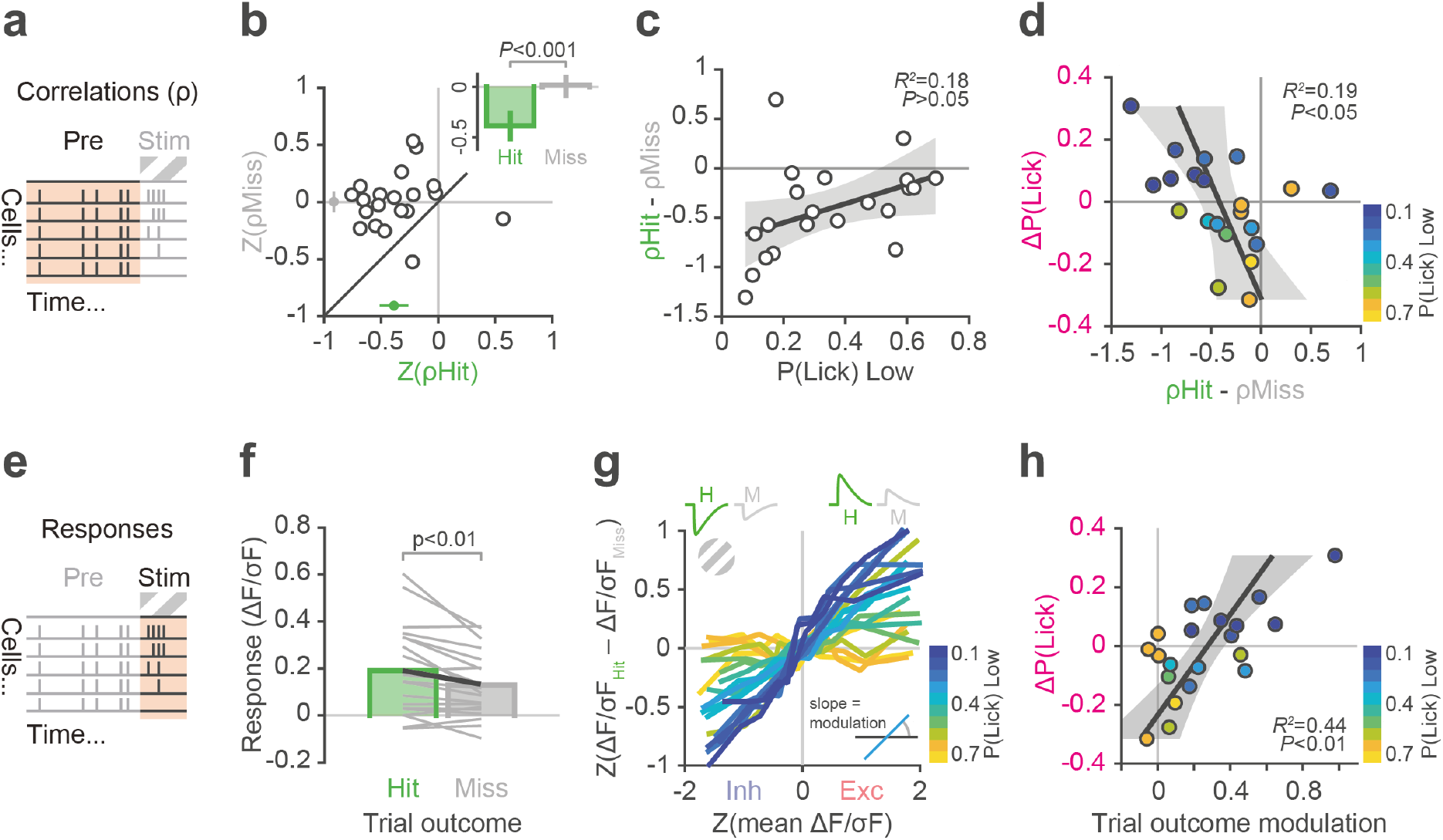
The state of cortex before and evoked by the visual stimulus influences the behavioural impact of photostimulation. **a.** Cartoon indicating the period of average spontaneous pairwise correlations preceding the initiation of the behavioural trial and presentation of the visual stimulus. **b.** The average pairwise spontaneous correlations between all cells within a session before hit trials are lower than before miss trials. Each data point is one experiment (*N* = 21 sessions, 14 mice), the green point indicates the mean correlation before hit trials, the grey point indicates mean correlation before miss trials (comparison also shown inset, error bars indicate 95% CI). **c.** The difference in pre-stimulus network correlations between hit and miss low-contrast trials (resampled 10,000 times to match hit and miss trial numbers) within a session is plotted against the overall task performance on low-contrast trials in that session. Each data point is one session. **d.** The photostimulation-mediated change in detection rate of the low-contrast stimulus (ΔP(Lick)) in a session is plotted against the difference in the pre-stimulus average network correlation between hit and miss low-contrast trials on that session. Each data point is one session and is coloured by the animal’s performance on low-contrast trials without photostimulation. **e.** Cartoon indicating behavioural trial and visual stimulus evoked responses. **f.** There is more activity evoked on average in visually-responsive cells on hit trials than miss trials. Each grey line is one session. **g.** The trial-outcome modulation of evoked response magnitude depends on overall task performance. For all cells in each experiment, the difference in evoked response on hit and miss trials (y-axis) is binned (10 equal sized bins) by average response magnitude (x-axis) of that cell regardless of outcome (resampled 10,000 times to match hit and miss trial numbers). The trial outcome modulation of the entire recorded population in a given session is then defined as the linear slope of the binned response versus outcome-modulation relationship. Each line represents one session, coloured by performance on low-contrast trials in that session. **h.** The slope of modulation of responses by trial outcome for a given session, which we interpret as a sign of active cortical engagement, correlates with the behavioural effect of photostimulation (ΔP(Lick)) in that session. Each point is one session, coloured by performance on low-contrast trials without photostimulation.

That pre-trial network synchronisation is a better predictor of trial outcome in low performance conditions (compared to high-performance conditions) suggests that cortex was more actively engaged and played a more prominent role in solving the task. This interpretation was confirmed on photostimulation trials where we found a significant relationship between the difference in network synchronicity before hit and miss trials, and the effect of photostimulation on behaviour (**Fig. 2d**, *R^2^* = 0.19, *P* = 0.048. *N* = 21 sessions, 14 mice. See also **Supplementary Fig. 3**). In summary, when animals performed poorly at detecting the low-contrast stimulus, the baseline correlation structure of the cortical network was a better predictor of performance, and in this behavioural state photostimulation of CT ensembles improved the detection of stimuli.

Next, we examined the relationship between activity evoked by the behavioural trial and task performance, reasoning that activity evoked by the low-contrast stimulus is likely to be a stronger determinant of its perceptual salience than background activity before the stimulus. We first looked at the activity resulting from the behavioural trials with low-contrast visual stimuli without photostimulation (**Fig. 2e**). On average, low contrast hit trials were associated with more evoked activity in neurons that were responsive to high-contrast visual stimuli (**Fig. 2f.** Hit: 0.19 ± 0.19 ΔF/σF, Miss: 0.13 ± 0.13 ΔF/σF measured in a 500 ms window immediately after the end of stimulus presentation. *P* = 0.0041 Wilcoxon signed rank test. *N* = 21 sessions, 14 mice). We looked into this further by examining how behavioural outcome is encoded in the activity of all recorded neurons. We investigated how the modulation of evoked activity by trial outcome depended on the mean visual response of a neuron and the overall performance level of the mouse in the session during which each neuron was recorded. More strongly visually responsive neurons showed stronger modulation by behavioural output such that neurons excited by the visual stimulus on average were relatively more excited on hit trials in comparison to misses, and neurons suppressed by the stimulus were more suppressed, producing a positive relationship between the mean response of neurons to the visual stimulus and the extent of their modulation by trial outcome (**Fig. 2g**). Furthermore, we found that the slope of this relationship, which defines the extent to which a population is modulated by trial outcome, depended on the overall performance on the session in which the neurons were recorded (**Fig. 2g.** and **Supplementary Fig. 4 and 5**). In sessions with good overall performance there was little to no modulation of evoked activity by trial outcome, but when performance was poor there was a large difference between the activity evoked on hit and miss trials. Similar to our findings on pre-trial correlation structure, when animals found the task challenging, cortical activity was more heavily modulated by trial outcome.

These results suggest that the relationship between cortical activity and performance depends on perceptual demand. In other words, when the detection of low contrast stimuli is perceptually demanding for the mouse, V1 activity is more tightly linked to behavioural performance, displaying less correlated activity in general (**Fig. 2c**) and a larger dynamic range of activity evoked by the stimulus (**Fig. 2g**). This indicates that these activity states reflect an active role of V1 when the perceptual demand is high. We next tested this hypothesis by examining photostimulation trials and asking how the cortical signatures of active engagement correlate with the influence of photostimulation on behaviour. We found a positive relationship between the effect of photostimulation on stimulus detectability and the extent of trial outcome modulation of the recorded population in the same sessions (**Fig. 2h.** *R^2^* = 0.44, *P* = 0.0011; *N* = 21 sessions, 14 mice). When trial outcome modulation was large, the effect of photostimulation was to increase stimulus detection and this corresponded to sessions where animal performed relatively poorly. Conversely, when mice performed the task easily, V1 was more passive in its representation of stimuli and associated trial outcome and photostimulation under these conditions suppressed the detection of visual stimuli. Together, this confirms a shift in the role served by cortical activity (from beneficial to suppressive) depending on perceptual demand.

How does photostimulation influence behaviour? The answer will depend on how activity propagates from the directly stimulated neurons. To begin answering this question we analysed how targeted photostimulation engaged the local circuitry. We first examined the patterns of network activity evoked by photostimulation of neuronal ensembles in the absence of a visual stimulus (**Fig. 3a**) and found that responsive neurons in the local network could be either excited or inhibited (**Fig. 3b**). The dominant effect of photostimulation was inhibition of other pyramidal cells revealing the known pattern of dense inhibitory connectivity ^55–61^. As we increased the number of photostimulated cells, the number of inhibited cells in the local network scaled approximately linearly with the number of activated cells (**Fig. 3c**, for target-zone neurons: *R^2^* = 0.71, *P* < 0.001; excited network neurons: *R^2^* = 0.18, *P* < 0.001; inhibited network neurons: *R^2^* = 0.33, *P* < 0.001; *N* = 14 mice, 63 sessions (stimulation ensembles pooled)). Concomitant with the increased inhibition we saw a reduction of the number of spontaneously excited cells. The progressive recruitment of inhibition resulted in the overall activity levels in the network remaining approximately constant as the number of photostimulated cells increased (**Supplementary Fig. 6**). We next examined the spatial distribution of activated and inhibited cells by creating a photostimulation-triggered spatial average of responding cells. The excited cells were localised in a narrow zone around each targeted cell, whereas the inhibited cells had a more widespread distribution, forming an annulus around the directly targeted cells (**Fig. 3d**). The differences between these distributions produces an overall centre-surround motif of excitation and inhibition, where the spatial spread of inhibition is larger than that of excitation (**Fig. 3e**, spread of excitation: 89 ± 48 μm; inhibition: 166 ± 51 μm. *P* < 0.001 Wilcoxon signed rank test. *N* = 14 mice, 63 sessions (stimulation ensembles pooled; see also ^50^). Interestingly, we found no differences in the total evoked activity or the spatial profile amongst the three different classes of stimulation ensemble (**Supplementary Fig. 6**).

**Figure 3.**
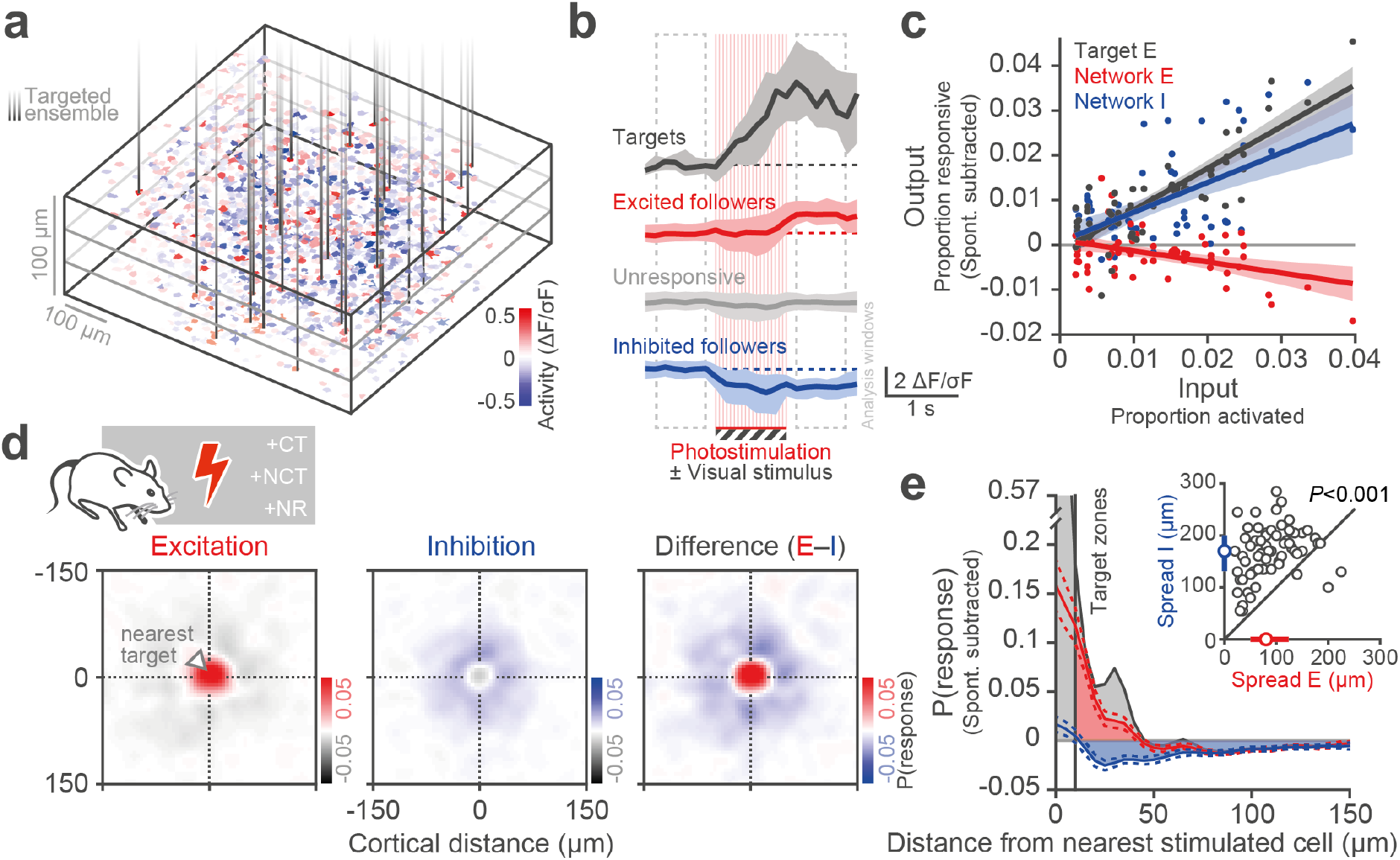
Photostimulation reveals net-inhibitory and centre-surround circuit architecture. **a.** Example segmented ROIs from one FOV coloured by evoked response to photostimulation during grey screen periods (catch trials). 4 planes were imaged simultaneously. Vertical black lines indicate SLM targets (35 cells targeted for simultaneous activation in this experiment). **b.** Example responses from a single trial showing the averages responses in directly targeted cells (black), detected excited responders (red, mean fluorescence in the response window is greater than 1 SD of the baseline window), non-responders (grey), and inhibited responders (blue, mean fluorescence in the response window is less than −1 SD of the baseline window). Bold lines indicate the median and shading indicates the interquartile range across all detected cells on this trial. **c.** Input-output function of the local network. The probability of detecting an excitatory (red) or inhibitory (blue) response in the network (excluding directly targeted cells) across all trials without visual stimulus, after subtracting the spontaneous rate, is plotted as a function of the proportion of cells stimulated. Each point represents the average response in one session, coloured by the type of response being measured (Target cell excitation (black), non-targeted network excitation (red) or inhibition (blue)). The lines indicate the fit, of recorded response against proportion of cells photostimulated, through all sessions. The shading indicates the 95% CI of the fits. **d.** The spatial profile of photostimulation is revealed by plotting the probability of detected responses (within the same plane as the stimulated cells) in 10×10 μm spatial bins, where each non-stimulated cell is positioned relative to its nearest stimulated target cell. Left: The spatial profile of the probability of detecting excitatory responses in the same plane as the stimulated cells after subtracting the probability of response seen in spontaneous periods. The spatial profiles are gaussian blurred (sigma = 10 μm) within each session and averaged across all sessions (*N* = 21 sessions, 14 mice). Middle: The probability of detecting inhibitory responses. Right: The difference between the excitatory and inhibitory response profiles reveals a small focal region of net excitation surrounded by an annulus of net inhibition. **e.** Quantification of the mean collapsed spatial profile of response probability similar to **d.** but across all recorded planes. Directly targeted cells and excluded nearby cells shown in black (the peak seen at 33 μm are likely indirectly stimulated cells immediately above and below a directly targeted cell), excitation of non-targets shown in red, and inhibition shown in blue. The dashed lines indicate 95% CI. Inset: The functional spread of inhibitory responses is wider than the spread of excitatory responses. The marginal coloured dots indicate the median and interquartile range.

These findings suggest that the effect of stimulating specific ensembles on behaviour (**Fig. 1**) cannot be explained simply by their influence on the overall activity of the local circuit. We therefore examined the effect of photostimulation on the functional identity of the recruited network and how that modified the neural representation of the visual stimulus, given that preferential connections between similarly tuned neurons have been reported in V1^55,62,63^. We thus restricted our next analysis to visually responsive neurons. We plot the probability of response of a neuron to the low contrast visual stimulus with and without photostimulation as a function of two variables: first, the tuning similarity to the fixed orientation of the low contrast visual stimulus, and second, the physical distance between the neuron and the nearest photostimulated neuron (**Fig. 4a**). Photostimulating CT ensembles during the visual stimulus enhanced the responses of nearby (< 50 μm) cells with tuning matched to the stimulus and therefore, the CT ensemble. The responses of dissimilar cells at these distances were suppressed. Outside of this focal zone (> 50 μm) we observed net and indiscriminate suppression of visually evoked responses. This motif of selective enhancement and suppression of cells in the local circuit when stimulating CT ensembles sharpens the population tuning curve of the network (**Fig. 4bc.** Population OSI change for neurons within 50 μm of the nearest stimulated cell when stimulating CT: 0.26 ± 0.47; NCT: −0.02 ± 0.41; NR: −0.14 ± 0.40. CT versus NCT *P* = 0.0045, CT versus NR *P* = 0.0078, NCT versus NR *P* = 0.1672. For neurons greater than 50 μm away from the nearest stimulated target when stimulating CT: 0.05 ± 0.09; NCT: −0.03 ± 0.23; NR: 0.00 ± 0.21. CT versus NCT *P* = 0.0129, CT versus NR *P* = 0.1808, NCT versus NR *P* = 0.3754. Wilcoxon signed rank test with Bonferroni multiple comparison correction). The sharpening of the population tuning curve was a specific result of stimulating cotuned, visually-responsive cells (**Supplementary Fig. 5**). Moreover, stimulation of non-visually-responsive neurons inhibited the visually-responsive neurons (**Fig. 4b**, **Supplementary Fig. 7**), revealing the existence of multiple competing populations.

**Figure 4.**
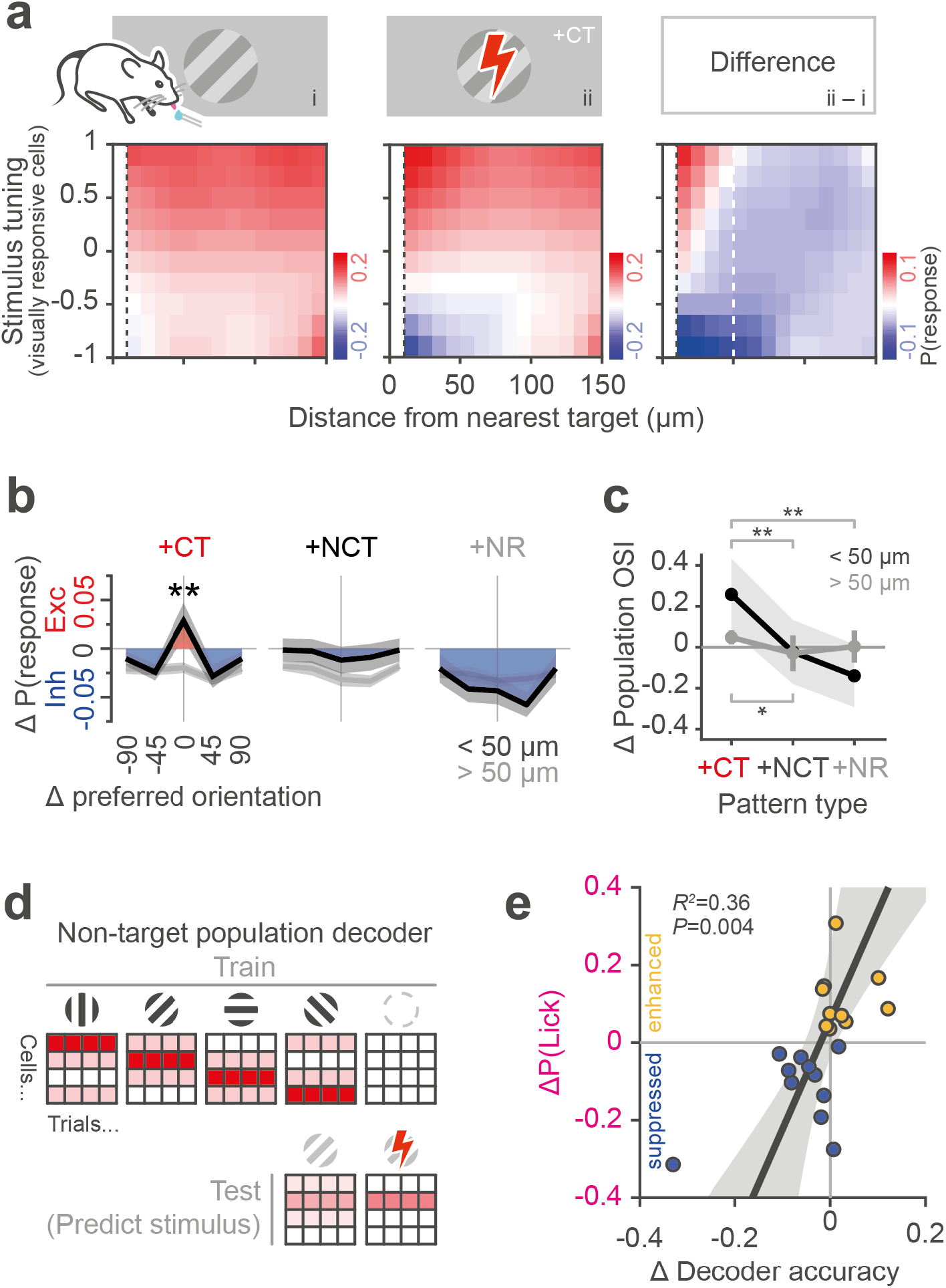
Photostimulation reveals like-to-like circuit architecture which impacts the stimulus encoding capacity of the network and the behavioural report of the stimulus. **a.** Restricting analysis to the non-stimulated visually-responsive subpopulation of neurons to relate sensory stimulus tuning to connectivity. Left: The probability of response of neurons to low contrast visual stimulus, binned by selectivity to the visual stimulus and by distance to the nearest stimulated target cell (note, there is no photostimulation in these trials), averaged across all experiments (*N* = 21 sessions, 14 mice). Middle: Responses to low contrast visual stimulus but with simultaneous photostimulation of CT ensemble. Right: The subtraction of visual-only from visual with photostimulation reveals that photostimulation causes net inhibition with feature-specific excitation and inhibition in a small spatial region close to the stimulated cells. Convolved with a gaussian filter, sigma = 1 bin (10 μm) for display only (*N* = 21 sessions, 14 mice). **b.** Population orientation tuning curves. The photostimulation evoked change in the visual stimulus response for all visually-responsive cells, binned by orientation preference (aligned to the stimulus orientation used for the experiment; Δ0 is the orientation of the visual stimulus and thus the preference of the cotuned ensemble) and then averaged across experiments. Two curves are shown, one for cells within 50 μm of the target cells (black) and one for cells further than 50 μm away from the nearest target cell (grey). Data at Δ-90 is the same as Δ+90, for display only. Thick lines show the mean and the shaded errors bars show the standard deviation. **c.** The selectivity (ratio of the value at Δ0 compared to the baseline, termed OSI) of the change in population orientation tuning curves from **b.** when stimulating each of the 3 types of ensemble. Thick lines indicate the mean across animals and sessions and error bars indicate the 95% confidence intervals. **d.** A decoder was trained to classify stimulus presence and orientation given the activity of visually-responsive non-target cells on high contrast trials. The classifier was then tested on low contrast trials with and without simultaneous photostimulation. **e.** The session average behavioural change of an animal detecting the low-contrast visual stimulus, caused by photostimulation (ΔP(Lick)), is correlated with the photostimulation-mediated change in the accuracy of stimulus decoding from population activity.

To relate these circuit level effects of photostimulation to behaviour we trained a classifier to decode the presence and orientation of a visual stimulus from activity of non-targeted cells on high-contrast trials. We then tested the decoder on activity evoked by the low-contrast visual stimulus with and without photostimulation (**Fig. 4d**). As expected, when the low-contrast evoked activity pattern more closely resembled that evoked by the high-contrast stimuli the decoder performed better (**Supplementary Fig. 8**). Overall decoder performance did not correlate with animal performance across all sessions, with no difference in performance seen between hit and miss trials (**Supplementary Fig. 8**). However, a strong relationship was observed between the photostimulation-mediated change in decoder performance and the associated change in behavioural detection of the stimulus (**Fig. 4e**, *R^2^* = 0.36, *P* = 0.004). When photostimulation acted to improve the stimulus-decoder performance the animal’s perceptual performance also improved. Conversely, when the stimulus representation was impaired by photostimulation the perceptual detectability of the stimulus was suppressed. These results thus provide a causal link between neuronal stimulus encoding and behavioural performance.

Our results provide a new perspective for understanding the link between activity in sensory cortex and perception, and for interpreting the consequences of perturbation experiments. We demonstrate that the behavioural effect of activation of cortical ensembles depends on their functional identity, with ensembles that normally represent the stimulus having the most potent effect. However, the effect of stimulating these ensembles has a bidirectional effect on behaviour: either boosting or inhibiting detection behaviour depending on task difficulty for the animal. Furthermore, we show that stimulating appropriate ensembles recruits postsynaptic activity in a functionally specific manner. This is likely a consequence of the wiring specificity of the local circuit ^50,55,62,63^. Functionally specific postsynaptic recruitment would be expected to alter stimulus representation and potentially behaviour. Consistent with this, we find that the success of a linear decoder of stimulus identity is altered by photostimulation and changes in decoding success are correlated with changes in behaviour in response to photostimulation.

To examine the interplay between these factors we constructed a multiple linear regression model (**Fig. 5a, b.** Full model *R^2^* = 0.75, *P* < 0.001), to define the relative influence of different neuronal population activity parameters in determining behavioural outcome in response to photostimulation. The model revealed that the dominant effects are the state of cortex as measured by the trial-outcome modulation of visually evoked responses (**Fig 2h**, referred to as ‘State_Stim_’), and the photostimulation-induced modification to the stimulus encoding of the network (**Fig. 4e**, referred to as ‘Activity_ΔStim_’) (**Fig. 5c**, correlation coefficients against the residual from otherwise complete models (semi-partial correlation) for State_Stim_: 0.51 [95% CI: 0.10 0.77], *P* = 0.017; Activity_ΔStim_: 0.52 [95% CI: 0.12 0.78], *P* = 0.014). The dual effects of cortical engagement and the stimulus decoding work in concert to explain the behavioural change caused by photostimulation. Despite being a simple linear model, the combination of neuronal population measures has strong predictive power for the influence of photostimulation on this detection behaviour (**Fig. 5d**, Reduced model, without State_Pre_, predicted values compared to actual values *R^2^* = 0.6, *P* < 0.001, RMSE = 0.08).

**Figure 5.**
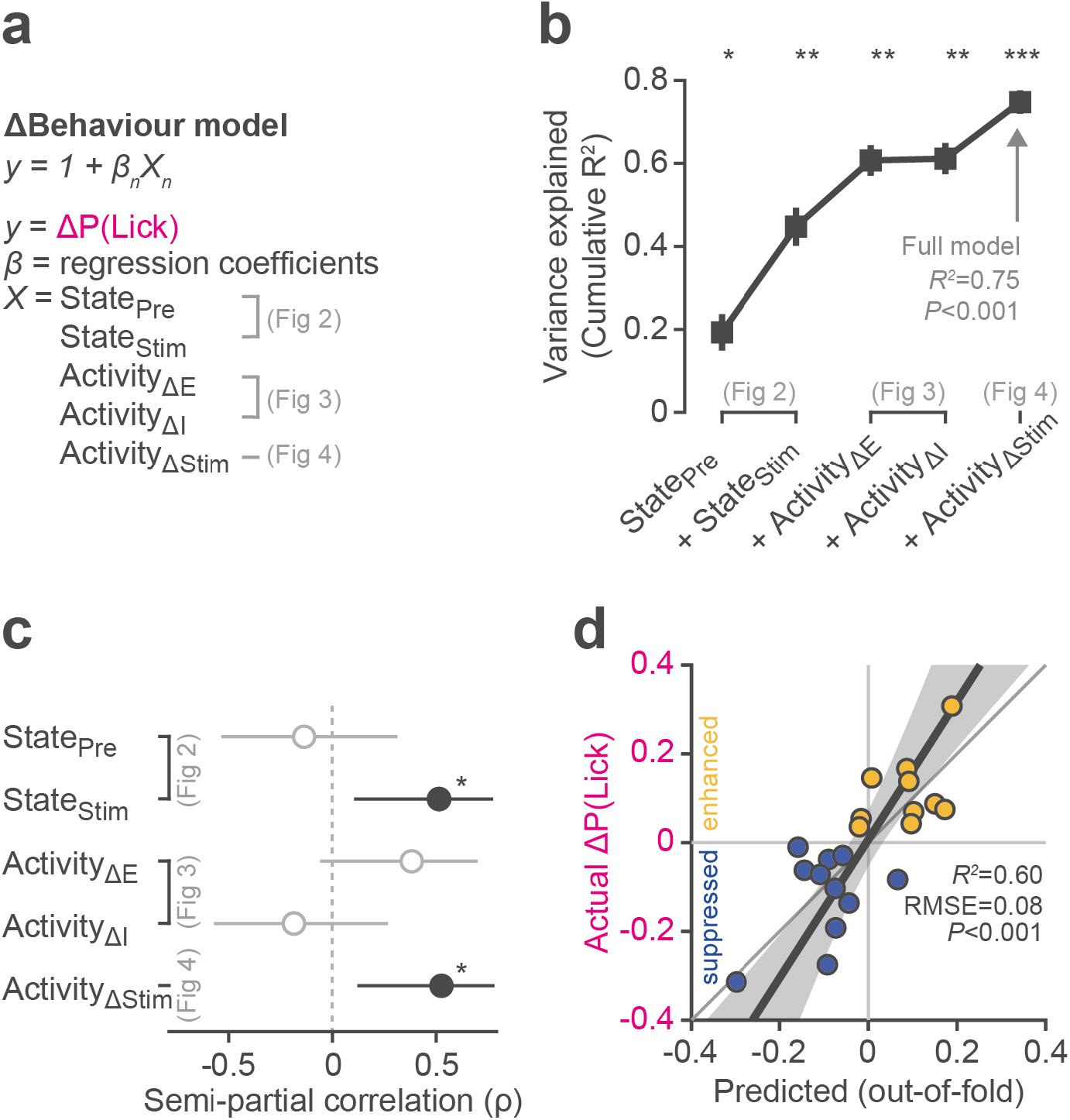
The behavioural effect of photostimulation is explained by both the cortical state of the animal and the evoked changes to local network activity. **a.** Multiple regression model for the session average effect of photostimulation on the change in detection behaviour, incorporating terms relating to the state of the animal (extracted from neural data pre and post stimulus (State_Pre_, State_Stim_), see **Fig. 2**) and the photostimulation-evoked change in local network activity (excitation/inhibition of all cells including targets (Activity_ΔE/I_), and the change in encoding capacity of the visually responsive cells (Activity_ΔStim_), see **Fig. 3**). **b.** Sequentially adding terms from results shown in previous figures to construct the full model. Errors bars indicate S.D. of the variance explained across all leave-one-out permutations (*N* = 21 sessions). **c.** Semi-partial correlations (the correlation between one model term and the residuals from a model containing all but that model term) of model coefficients against ΔP(Lick) show the relationship of the variables of interest while accounting for all other variables in the full model. Both the trial outcome modulation of responses (State_Stim_) and the change in stimulus decoding (Activity_ΔStim_) significantly explain the behavioural effect of photostimulation. Error bars indicate the 95% CI of the semi-partial correlation coefficient. **d.** The predictive capacity of the reduced (excluding State_Pre_) model is evaluated with leave-one-out cross validation by comparing the experimentally obtained ΔP(Lick) with the value predicted from the model excluding this session from the training set. The shaded region indicates the CI of the fit.

These results help to reconcile apparently contradictory findings about the engagement of the cortex in behavioural tasks^20,21,25–27^. Our results suggest that when task demands are high, primary sensory cortex is engaged and plays a positive causal role in determining task performance. When the task is easy to solve then alternative pathways and downstream networks are likely already optimally engaged, and cortical stimulation represents a distractor. Our results parallel recent findings from lesion and silencing experiments^26^ showing that learned tasks are no longer cortically dependent and additionally suggest that cortical resources are dynamically allocated depending on task difficulty and performance. Our findings differ from a recent study^64^ which found that photostimulation at just two cell locations could recruit an associated ensemble which then always positively biased stimulus perception. However, the perceptual demands of their discrimination task are greater than for our detection task, which is consistent with the interpretation of our results.

A prominent feature of the local network response to photostimulation was inhibition of other cells, similar to other reports^50,65,66^. This suggests that there exists strict control over the balance between excitatory and inhibitory activity levels^67^, similar to what has been observed during spontaneous and stimulus evoked activity states in awake visual cortex^68^. We did not observe large scale ‘pattern completion’ modes of activity triggered by photostimulation of cotuned cells as recently reported^64^ but rather observed a more constrained and balanced interplay between the identity and the activity of groups of cells. A recent study^50^ found that photostimulating single cells predominantly inhibited other similarly tuned cells, while excitation was reserved for a very small population of very similar cells. We hypothesise that the more relaxed relationship of influence versus stimulus tuning that we observed could arise through multiple cells being photostimulated coincidently in our experiments.

Our findings raise a number of outstanding questions. First, which downstream networks^69–73^, either cortical or subcortical, are charged with reading out the task-dependent information carried by the stimulated layer 2/3 neurons? This will require recording and manipulation of deeper cortical layers and subcortical target areas, as well as recording from multiple areas simultaneously. Second, which circuits, including neuromodulatory pathways^28,31,74^, are responsible for modulating the contribution of cortex to the task? Our results also raise a more general question of how and where multiple streams, including the collicular pathway^75,76^, of visual processing combine and interact. Finally, uncovering the detailed elements of the neural code underlying perception will require further refinements in the temporal precision, spatial resolution and physical coverage of both recording and stimulation approaches, as well as performing flexible real-time activity-guided manipulations^77^ in concert with sophisticated analytical frameworks^78^.

## Methods

All experimental procedures were carried out under license from the UK Home Office in accordance with the UK Animals (Scientific Procedures) Act (1986).

### Animal preparation

We used transgenic GCaMP6s mice (Emx1-Cre;CaMKIIa-tTA;Ai94 ^47^ and CaMKIIa-tTA;tetO-G6s ^48^) of both sexes aged between P41 and P73. Doxycycline treatment in drinking water from birth to P49 prevented interictal activity in the Ai94 mouse line ^79^. Briefly, to prepare the mice for all-optical experiments we excised the scalp and implanted a metal headplate. We then removed the skull and dura overlying visual cortex, injected virus encoding the opsin and implanted a chronic cranial imaging window in place of the skull. Sterile procedures were used throughout. Before surgery, mice were given a subcutaneous injection of 0.3 mg/mL buprenorphine hydrochloride (Vetergesic) and anaesthetised with isoflurane (5% for induction, 1.5% for maintenance). The scalp above the dorsal surface of the skull was removed and an aluminium headplate with a 7 mm diameter circular imaging well was fixed to the skull centred over the right monocular primary visual cortex (2.5 mm lateral and 0.5 mm anterior from lambda) using dental cement. A 4 mm diameter craniotomy was drilled inside the well of the headplate, and the dura was then carefully removed. A calibrated pipette bevelled to a sharp point (inner diameter ~15 μm) connected to a hydraulic injection system (Harvard apparatus) was used to inject small volumes of virus (AAV2/9-CaMKII-C1V1(t/t)-mRuby2-Kv2.1). The dilution of virus in buffer solution (20 mM Tris, pH 8.0, 140 mM NaCl, 0.001% Pluronic F-68) was adjusted throughout experiments to optimise expression levels and ranged from 8-fold to 25-fold dilution of stock (stock concentration: ~6.9×10^14^ gc/ml). We made ~5 insertions of the injection pipette, each site spaced by ~300 μm. At each site we slowly lowered the pipette to a depth of 300 μm below pia and injected 150 nl of the virus solution at 50 nl/min. After each injection the pipette was left in place for a further 3 minutes before slowly retracting. We then press-fit a chronic window (a 3 mm coverslip bonded to the underside of a 4 mm coverslip with UV-cured optical cement, NOR-61, Norland Optical Adhesive) into the craniotomy, sealed with cyanoacrylate (Vetbond) and fixed in place with dental cement (SuperBond). Following surgery, animals were monitored and allowed to recover for at least 7 days. After recovery we began behavioural training. All-optical experiments were then performed > 3 weeks post-surgery allowing for sufficient expression levels (animals were aged P67 – P134, median = P97 at time of experiments).

### Behavioural training

We used an operant conditioning protocol whereby headfixed mice were required to lick at a water spout positioned in front of them to report detection of a visual stimulus. Licks were recorded electrically. If the mice reported presence of the stimulus correctly a sugar water reward (10 % w/v sucrose) was delivered through the water spout. The behaviour hardware was controlled by custom software (PyBehaviour, https://github.com/llerussell/PyBehaviour) interfacing with an Arduino to trigger stimuli, record licks and deliver rewards. Mice had free access to food in their home cage but access to water was limited to that acquired during the task. Mice had their weight monitored before and after daily training and were supplemented with additional water to maintain a minimum of 80% of their starting body weight. Before training mice were habituated to handling and head restraint over 2 days. Training then took place in individual sound-dampened enclosures in which the mice were head-fixed and allowed to run on a treadmill. While not an integral part of the task design we found that allowing mice to run improved their performance in the task. Trials were triggered after mice withheld licks for 4 ± 3 seconds, after which a monocular visual stimulus appeared in the centre of the monitor. If the mice licked at the water spout at any point during the stimulus a reward was delivered. In the first few days of training a reward was delivered automatically at 800 ms. Mice quickly learnt the requirements of the task and their reaction times preceded this automatic reward delivery time. After a few days the automatic reward delivery was disabled. After the stimulus and response window there was a fixed inter-trial period of 7 seconds before the next ‘withhold’ period was started. We also delivered randomly interleaved catch trials (no visual stimulus) to record chance rate of licking and assess accuracy in the task. Once stable performance was reached, we progressed the mice to a psychophysical variant of the task where we introduced a range of contrasts to assess their perceptual threshold. We found that task performance was insensitive to stimulus location on the monitor. For the final experiment the trial order was pseudo randomised so as to ensure a constant rate of ‘easy’ trials and rewards while also ensuring that repeats of the same probe types were not immediately consecutive.

### Visual stimulation

Visual stimuli were generated using custom software (using PsychoPy ^80^). 30 ^o^ Gabor patches of drifting sinusoidal gratings (8 directions, 0 to 315 ^o^ in 45 ^o^ increments) with a spatial frequency of 0.04 cycles/^o^ and a temporal frequency of 2 Hz were presented on a monitor (typically 51.8 cm width, 32.4 cm height, 15 cm from the animals left eye covering up to ± 47 ^o^ of the vertical visual field and ± 60 ^o^ of the horizontal visual field), with a spherical distortion applied to correct perspective errors.

#### Training

During training the orientation of the stimulus was randomised on every trial and the duration of the stimulus was 1 second. Rewards were delivered if the mouse licked during the stimulus regardless of the orientation.

#### Mapping orientation preference

To map orientation preference with two-photon imaging the gratings were positioned in the retinotopically appropriate location and were presented in a randomised order with a duration of 3 seconds, interleaved by 5 seconds of mean luminance grey. If mice licked at the water spout during this mapping block a water reward was delivered.

#### Experiment

During the behavioural experiments with photostimulation the visual stimuli parameters were the same as during training except the stimulus was positioned in the retinotopically appropriate location for the imaging field of view. Two contrasts were used, high (100%) and low (range: 1 – 10%, mean ± SD = 4.8 ± 3 %). The direction of the high contrast stimulus was randomised on every trial. The direction of the low contrast stimulus was fixed to match the orientation preference of the ‘cotuned’ photostimulation ensemble.

### Widefield imaging

To locate primary visual cortex, and position the experimental field of view, widefield GCaMP imaging was performed (usable FOV ~ 2 × 2 mm). GCaMP6s fluorescence produced by one-photon excitation (470 nm LED, Thorlabs) was collected through a 5x/0.1-NA air objective (Olympus) onto a CMOS camera (Hamamatsu ORCA Flash 4.0, binned image size of 512 × 512 pixels, 20 Hz frame rate). Contrast-reversing checkerboard bars, 10° wide were drifted vertically and horizontally across a grey screen at a speed of 25 °/s in an interleaved sequence. Stimulus triggered change in fluorescence for the two different stimuli revealed areal borders and identification of primary visual cortex ^81^. This was repeated with two-photon imaging on the day of the experiment to confirm the retinotopic location of the chosen field of view.

### Two-photon population imaging

Two-photon imaging was performed with a resonant scanning microscope (Ultima II, Bruker Corporation) using a Chameleon Ultra II laser (Coherent) driven by PrairieView. A 16x/0.8-NA water-immersion objective (Nikon) was used for all experiments. An ETL (Optotune EL-10-30-TC, Gardasoft driver) was used to perform volumetric imaging, spanning a 100 μm range with 33.3 μm spacing between planes. The FOV size ranged from 600 × 600 to 850 × 850 μm, at a constant image size of 512 × 512 pixels. The number of cells recorded (ROIs) per experiment ranged from 316 to 4,454 (mean = 2,266 ± 1,607). In single-plane experiments the frame rate was 30 Hz, in volumetric experiments the per-plane frame rate was 7 Hz. GCaMP6s was imaged at 920 nm and mRuby (conjugated to C1V1-Kv2.1) was imaged at 765 nm. Power on sample was 50 mW at the shallowest plane (~150-200 μm below pia) and increased to ~85 mW at the deepest plane (~250-300 μm), interpolating for intermediate planes, to equalise imaging quality across planes. To maximise imaging quality ^82^ we calculated the tilt of the sample relative to the microscope and then rotated the objective along two axes to be perpendicular to the implanted coverslip window.

### Two-photon photostimulation

Two-photon photostimulation was carried out using a fibre laser at 1030 nm (Satsuma, Amplitude Systèmes, 2 MHz rep rate). The laser beam was split via a reflective spatial light modulator (SLM) (7.68 × 7.68 mm active area, 512 × 512 pixels, OverDrive Plus SLM, Meadowlark Optics/Boulder Nonlinear Systems) which was installed in-line of the photostimulation path (Neuralight, Bruker Corporation). Phase masks used to generate focused beamlet patterns in the sample were calculated via the weighted Gerchberg-Saxton algorithm. The targets were weighted according to their location relative to the centre of the SLM’s addressable FOV to compensate for the decrease in diffraction efficiency when directing beamlets to peripheral positions. We calibrated the targeting of SLM spots in imaging space by burning arbitrary patterns with the SLM using the photostimulation laser in a fluorescent plastic slide before taking a volumetric stack of the sample with the imaging laser. We manually located the burnt spots and the corresponding affine transformation from SLM space to imaging space was computed. For 3D stimulation patterns we interpolated the transformation required from the nearest calibrated planes (Calibration code: https://github.com/llerussell/SLMTransformMaker3D). To increase stimulation efficiency, we offset the photostimulation FOV with the photostimulation galvanometers such that the centre of SLM space was close to the cortical/imaging-space centroid of targeted cells. Spiral photostimulation patterns (3 rotations, 10 μm diameter, 20 ms duration) were generated by moving all beamlets simultaneously with the galvanometer mirrors. The laser power was adjusted to maintain 6 mW per target cell.

### Naparm (Near automatic photoactivation response mapping)

To find photostimulation-responsive cells we semi-automatically detected cell locations from expression images and stimulus-triggered average or pixel-correlation images (STA Movie Maker, https://github.com/llerussell/STAMovieMaker). These cell body coordinates were then clustered into equal size groups of user-determined size (between 10 and 50) and the groups were stimulated one by one. The associated phase mask, galvanometer positioning and Pockels cell control protocol were generated with custom MATLAB software (Naparm, https://github.com/llerussell/Naparm) and executed by the photostimulation modules of the microscope software (PrairieView, Bruker Corporation) and the SLM control software (Blink, Meadowlark). For responsivity mapping purposes we used a stimulation rate of 20 Hz, for 500 ms per pattern, and performed 8-10 trials. These data were then analysed online together with the visual response mapping data to extract activity traces and design stimulation ensembles (see below).

### Synchronisation

For subsequent synchronisation during analysis, analogue signals of various trigger lines were recorded with a National Instruments DAQ card, controlled by PackIO ^83^. The recorded inputs included two-photon imaging frame pulses, photostimulation triggers, galvanometer command signals, triggers to and frame flip pulses from the visual stimulus and the SLM phase mask update. Photostimulation trials for the responsivity mapping block were triggered at a fixed rate from an output line on the DAQ card. For the online behaviour experiments photostimulation and visual trials were triggered through the behaviour software and hardware.

### Experimental protocol

On the day of the full experiment the following protocol was used. First, we located an expressing region of cortex and quickly mapped the corresponding retinotopic location with two-photon imaging. After determining where to position the visual stimulus on the monitor we then presented drifting gratings of 8 different orientations while performing two-photon imaging to map orientation preferences of the recorded cells. Rewards were delivered during the visual stimuli if the mouse licked. Next, we stimulated a large proportion of all cells in the recorded volume to find which ones were photostimulation-responsive. Finally, we designed photostimulation patterns for use in the behaviour experiment (see below). We then gave animals ~10 trials to warm up before estimating the perceptual threshold for that animal on that day, after which the main behavioural experiment began. We recorded in 20 minute blocks, manually correcting for any drift in imaging FOV.

### Neuronal response metric

To measure neuronal responses we extracted the mean fluorescence in a ~500 ms window (4 frames for 3D experiments, 15 frames for 2D experiments) starting immediately after the photostimulation ended (and/or visual stimulus to ensure comparable measurements) and subtracted the mean fluorescence in the ~1 second baseline (7 frames for 3D experiments, 30 frames for 2D experiments) before the onset of photostimulation (or visual stimulus). We divided the difference in the means by the standard deviation of the baseline window, to give a signal-to-noise ratio (ΔF/σF). If on a given trial, for a given cell, this value was greater than 1 the response was scored as excited, and if it was below −1 the response was scored as inhibited (**Supplementary Fig. 9**). We excluded all photostimulation frames because of the associated artefact contaminating the activity traces. The slow kinetics of GCaMP6s permit this, although the magnitude of response is underestimated. We additionally compute a net response probability for each cell as the difference between that cells probability of being excited and inhibited across all trials. For comparison across experiments we then subtracted either the visually evoked response probability, or for trials without visual stimulation, the probability of detecting a positive or negative response in catch trials.

### Online photostimulation ensemble design

To increase speed of data analysis immediately prior to the experiment we streamed the raw acquisition samples to custom software (PrairieLink, RawDataStream, https://github.com/llerussell/Bruker_PrairieLink). We used this raw stream to process the pixel samples, construct imaging frames and in a subset of experiments, perform online registration. Processing online allowed us to directly output to a custom file format making the data immediately available for analysis. Motion corrected movies were loaded into MATLAB (MathWorks) and traces were extracted from both the photostimulation and the visual stimulation movies, using the photostimulation targets as seed points around which circular ROIs were dilated. We subtracted a neuropil signal from the ROI signal before determining responsivity. We determined cells as photostimulation-responsive if their evoked response (in a ~500 ms window after stimulus offset) to their direct stimulation was > 30% ΔF/F on > 50% trials. We determined cells as visually-responsive by the same criteria (with response window of 2 seconds during the stimulus presentation), additionally specifying their preferred orientation as the stimulus that elicited the largest average response (**Supplementary Fig. 10**).

Three types of task-relevant stimulation patterns were then designed in each experiment. After filtering for photostimulation-responsive cells the groups were designed and matched for number of activated cells, average evoked response magnitude and spatial clustering (average pairwise distance and nearest neighbour distance) but differed maximally in sensory tuning. The cotuned group was selected first, taking the largest group of photostimulation-responsive and orientation tuned neurons (minimum number of targets: 4, maximum: 79, median: 17) and thus set the constraints for the other groups to match. Groups were matched within session, not across sessions (**Supplementary Fig. 11**).

### Pre-processing: Imaging frame registration, ROI segmentation and neuropil correction

For the final analysis the raw calcium imaging movies were pre-processed using Suite2p ^84^. The pipeline included image registration, segmentation of active region of interest (ROIs), and of local surrounding neuropil signal. The final selection of ROIs was filtered semi-automatically using anatomical criteria to include only neuronal somata and discard spurious ROIs. We manually inspected all FOVs to ensure consistent results. We subtracted a neuropil signal from every ROI signal. The contamination of the ROI signal by the neuropil signal depends on many factors including expression levels, imaging quality, and axial sectioning by the imaging plane. We used robust linear regression to estimate the coefficient of neuropil contamination for each ROI (**Supplementary Fig. 12 ^46^**). The slope of this fit was used to scale the neuropil signal before subtraction from the ROI signal, such that after subtraction there was no correlation between the ROI baseline and neuropil. Neuropil subtraction had minimal effect on the response magnitude and negative responses were seen even without subtracting the neuropil contamination (**Supplementary Fig. 13,14**).

### ROI exclusion zones

In order to reduce potential off-target photostimulation artefacts we excluded from consideration all cells within a 20 μm diameter cylinder extending through all axial planes when analysing the network response to photostimulation due to potential imaging and photostimulation artefacts (see **Supplementary Fig. 13**). We redefined our target stimulation pattern identities based on the ROIs segmented by Suite2p within the 20 μm lateral disk around each of the SLM target locations. We also excluded ROIs in the first 100 rows of pixels of each imaging frame due to an ETL artefact related to the settle time of the lens when changing planes.

### Behavioural session truncation

To ensure we only analysed periods of the behavioural session where the mice were similarly engaged and motivated, we truncated the session when the rolling average performance (20 trial sliding window) of the ‘easy’ high contrast trials dropped below 70% of the starting performance.

### Data exclusion criteria

We excluded trials if > 50% of photostimulation targets failed to respond on that trial. We also excluded trials if the mice licked early (within the first 150 ms of the presentation of the visual stimulus). Whole sessions were excluded if fewer than 10 trials in the low-contrast or low-contrast with photostimulation condition remained (the median minimum number of trials in included sessions = 31 trials (range 11 – 69)). Out of 30 completed sessions, 9 were excluded (5 because too few trials remained, and 4 because of poor photostimulation efficiency)

### Statistical procedures

No statistical methods were used to predetermine sample size. The experiments were not randomised, and investigators were not blinded to allocation during experiments and outcome assessment. Summary statistics in the text are reported as mean +/− SD unless otherwise indicated. Statistical tests used are specified in the text and were generally two-tailed and nonparametric.

### Behavioural effect of photostimulation resampling procedure

To assess statistical significance, we devised a procedure to determine whether the photostimulation induced perceptual bias we observed across sessions could occur by chance through behavioural variability. For each session we had a mean lick rate to low contrast stimuli (baseline performance), and a mean lick rate to low contrast stimuli with photostimulation. When we plot these values from every session against each other, we get a slope that deviates from 1. To assess the significance of this slope, we generated many resampled “fake” lick rates for each session, then took one resample per session for every session and calculated a “fake” slope across sessions for each resample. Each resample for each session was constructed by sampling the same number of trials as were available for that session in the low-contrast photostimulation trial type where each individual trial had the same probability of being a hit (lick) as the real mean lick rate to the low contrast stimuli. We did this 10,000 times and then asked if the real slope across all sessions fell outside of the resampled distribution of slopes.

### Pre-trial correlations

To compute the network synchrony prior to presentation of the visual stimulus we used deconvolved activity traces (OASIS ^84,85^) smoothed with a Gaussian filter (sigma = 0.5 s). We used a 4.5 – 0.5 s window immediately prior to the initiation of the trial (delivery of a stimulus, if not a catch trial) as the ‘pre-trial’ period. We then computed pairwise correlations within these windows and averaged together all pairwise correlation coefficients across all cells (including targets) to give the total network correlation. We then z-scored all network correlations within animal and across all trial types to facilitate across animal comparisons. When comparing hit and miss trials we resampled 10,000 times to match trial numbers.

### Stimulus decoder

We used a multiple-class support vector machine (SVM) to decode and classify trial type (presence and orientation of high contrast visual stimulus) within a session, in which the output of multiple binary classifiers are compared to one another. We only used ROIs which were determined to be ‘visually responsive’ and excluded all target and nearby ROIs. We randomly selected half of the high contrast visual stimulus trials and half of the catch trials (no visual stimulus) in a session to train that sessions’ model. The remaining 50% of trials were used for cross-validating the performance on the held out high contrast and catch trials. We repeated this cross-validation procedure 100 times. We evaluated the high-contrast models with all of the available low-contrast trials. Note there was only one orientation of low-contrast stimulus in each session. We averaged the test results across all 100 permutations of the trained models for each session.

### Code availability

Custom code used for data acquisition, photostimulation control, behavioural training and analysis have been deposited online:

Naparm (https://github.com/llerussell/Naparm) PyBehaviour (https://github.com/llerussell/PyBehaviour) 3D SLM calibration (https://github.com/llerussell/SLMTransformMaker3D) STAMovieMaker (https://github.com/llerussell/STAMovieMaker) RawDataStream (https://github.com/llerussell/Bruker_PrairieLink) Objective rotation (https://github.com/llerussell/MONPangle)

## Acknowledgments

We thank Matteo Carandini, Kenneth Harris and Arnd Roth for helpful discussions and comments on the manuscript; Soyon Chun and Agnieszka Jucht for mouse breeding; Maite Marcantoni and Florence Bui for behavioural training during pilot experiments; Arthur Gretton and Peter Latham for analysis advice; and Bruker Corporation for technical support. This work was supported by grants from the Wellcome Trust, Gatsby Charitable Foundation, ERC, MRC and the BBSRC.

## Author contributions

LER performed surgeries and performed all-optical experiments, and analysed all data. ZY and LPT trained animals. MF and AMP provided valuable advice about experimental design and analysis. LER built hardware and software for control of behavioural training and for calibration and control of photostimulation. HWPD helped build photostimulation control software. SC and CDH provided C1V1-Kv2.1 virus. LER and MH conceived and designed the study and wrote the manuscript with input from all authors.

