## Supplementary figures for "The influence of visual cortex on perception is modulated by behavioural state"

**Supplementary Figure 1. Task acquisition and contrast threshold**

a) Lick raster plots, sorted by trial type (go and catch) for the first, second and fifth day of behavioural training for one example animal. Black dots indicate the first lick (reaction time) after presentation of the visual stimulus. Blue line indicates the timing of the automatically delivered water reward (AR) in the early training sessions.
b) Performance improves across sessions for all mice included in this study. Performance is defined as the probability of the first lick occurring (before the automatic reward delivery if present).
c) Reaction time decreases as the mice learn the task.
d) Example lick raster for one mouse during a psychophysical assessment session. Trials were delivered in random order but are sorted by stimulus contrast for display.
e) Min-max normalised probability of detecting a lick during the stimulus presentation at different contrasts. Performance decreased as stimulus contrast decreased. The black circle indicates the average contrast threshold (50% detection rate at 3.0 ± 1.7 % stimulus contrast).
f) Average time of the first lick (reaction time) to the different stimulus contrasts, at any point in the trial after the stimulus presentation.

**Supplementary Figure 2. Behavioural effect of photostimulation for all three stimulation ensembles.**

a) Illustration of the resampling based statistical procedure to determine significance of the relationship between the stimulus detection rates (P(Lick)) on trials without and trials with photostimulation. For each session we generated a resampled session using the mean response rate and number of trials for trials without photostimulation. We computed the slope and intercept of the fit between the real response rate to the visual stimulus and the resampled response rate. We repeated this procedure 10,000 times. We then compared the slope and intercept obtained from the fit to experimental data – including photostimulation trials – to the distribution of slopes obtained through the resampling procedure.
b) The relationship of stimulus detection rates (P(Lick)) within a session on trials without and trials with CT ensemble photostimulation. P-values are computed from the percentile of the resampled distribution in which the real slope lies. The unity line is shown in grey. The shaded region indicates the 95% CI of the fit.
c) Same as for b) but for trials with NCT ensemble stimulation.
d) Same as for b) but for trials with NR ensemble stimulation.
e) Similar to b) the relationship between the stimulus detection rate without photostimulation and the change (ΔP(Lick)) caused by photostimulation of the CT ensemble.
f) Same as for e) but for trials with NCT ensemble stimulation.
g) Same as for e) but for trials with NR ensemble stimulation.

**Supplementary Figure 3. The relationship between pre-trial correlations and task performance.**

a) Plotting the z-scored pre-trial correlation on only hit trials against performance on the low-contrast trials reveals a relationship between the value of correlation on hit trials with overall performance. Right: When plotting against the change in performance caused by photostimulation a clear relationship emerges: Enhanced stimulus detection as a result of photostimulation is associated with sessions in which hit trials have a lower pre-trial correlation.
b) Same as for a) but for correlations before miss trials. No relationship between miss trial pre-trial correlation and behaviour is seen.

**Supplementary Figure 4. The trial-outcome modulation of evoked response magnitude depends on overall task performance.**

The slope of the recorded population stimulus response modulation by trial outcome is plotted against the overall performance in the low-contrast trials in the session in which it was recorded. Colour indicates the change in stimulus detectability on trials with photostimulation in these sessions.

**Supplementary Figure 5. Average responses of the most and least responsive sub-populations on behavioural trials with low contrast visual stimulation.**

a) The average response on hit and miss trials for the bottom 10% of all cells (sorted by average response magnitude) in each session. These are cells which are most inhibited by the visual stimulus on average. The diagonal unity line is shown in grey. Each dot represents the average population response in one session, coloured by the stimulus detection rate (P(Lick)) on low contrast visual stimulus trials.
b) Same as for a) but for the top 10% responsive cells in each session.
c) The average response of the bottom 10% of cells (same as in a)) on hits (green) and misses (grey) plotted against the behavioural performance in the low contrast visual stimulus trials.
d) Same as in c) but for the top 10% of all cells in each session. The biggest modulation of response occurs in hit trials of the low performance sessions.
e) The difference between average responses on hit and miss trials for the least (bottom 10%, blue) and the most (top 10%, red) responsive of all cells in each session, plotted against the behavioural performance.

**Supplementary Figure 6. Network response to photostimulation for all three stimulation ensembles.**

a) Left: The network output (difference in the probabilities of detecting an excitatory or an inhibitory response) in either just the target cell population (red), the entire recorded population including targets (black) or the entire recorded population excluding the target cells (blue) when photostimulating CT ensembles, plotted against the proportional number of cells stimulated (number of targets divided by number of all recorded cells). Right: The average excitatory, and inhibitory, and the difference (net), response probabilities, for subpopulations of the recorded network. Wilcoxon signed rank test. * denotes $P < 0.05$ after Bonferroni correction.
b) Same as for a) but for stimulation of NCT ensembles.
c) Same as for a) but for stimulation of NR ensembles.
d) The average net difference in network activity (detection of excitatory and inhibitory responses) when stimulating all three ensemble types in all cells including targets (left) or excluding targets (right). Wilcoxon signed rank test with Bonferroni correction.
e) Left: The spatial profile of the network (excluding target cells) response to photostimulation of all three ensemble types. Right: Quantification of the spread of excitatory and inhibitory responses.

**Supplementary Figure 7. Competition between visually responsive and non-responsive populations**

Splitting the recorded population into sub-groups of visually-responsive (excited) or non-responsive reveals competition between the sub-populations. Stimulating the non-visually responsive ensemble during visual stimulus presentation leads to more suppression of the visually-responsive population on average when compared to stimulation of the two visually responsive ensembles (CT and NCT). Wilcoxon signed rank test with Bonferroni correction. *N* = 21 sessions, 14 mice. * = $P < 0.05/15$, ** = $P < 0.01/15$, *** = $P < 0.001/15$.

**Supplementary Figure 8. Population activity visual stimulus decoder.**

We used a multiple-class support vector machine (SVM) to decode and classify trial type (presence and orientation of high contrast visual stimulus) within a session, in which the outputs of multiple binary classifiers are compared to one another. We only used ROIs which were determined to be 'visually responsive' and excluded all target and nearby ROIs. We randomly selected half of the high contrast visual stimulus trials and half of the catch trials (no visual stimulus) in a session to train that sessions' model. The remaining 50% of trials were used for cross-validating the performance on the held out high contrast and catch trials. We repeated this cross-validation procedure 100 times. We evaluated the high-contrast models with all of the available low-contrast trials. Note there was only one orientation of low-contrast stimulus in each session. We averaged the test results across all 100 permutations of the trained models for each session.

a) Illustration of the classifier comparison scheme. Shaded boxes indicate the unique comparisons being made. Light grey boxes indicate classifiers comparing activity evoked by any orientation of stimulus (labels 1:4) with catch trials (no stimulus, labelled 0) which we refer to as ‘detection’ classifiers. Dark gey boxes indicate classifiers comparing two orientations of stimulus and we refer to them as ‘discrimination’ classifiers.

b) Example of the trained weights assigned to individual ROIs in an example ‘detection’ classifier. Cells with a larger evoked response to visual stimuli are weighted towards the stimulus class. There is no relationship between the stimulus selectivity of ROIs with the class they are weighted towards.
c) Example of the trained weights assigned to individual ROIs in an example ‘discrimination’ classifier. Cells that respond more to the visual stimulus have a stronger weight assigned to them. There is a clear relationship between stimulus specificity (the difference in response to the two stimuli under consideration) and the sign of the classifier weight.
d) Cross validation performance when testing the trained classifiers on the 50% of held-out high contrast and catch trials. Averaged across 100 permutations. Decoder accuracy is the true positive rate – the proportion of times the decoder correctly assigns trial type identity. Note that because we exclude target cells and nearby cells (within a 20 μm diameter cylinder, extending through all axial planes) we construct 3 classifiers per session, one for each stimulation ensemble type where different cells were stimulated and thus excluded. The classifier for NR stimulation trials performs slightly but significantly better (Wilcoxon signed rank paired, two-sided test with Bonferroni correction. * = $P < 0.05/3$, ** = $P < 0.01/3$, *** = $P < 0.001/3$). This is likely because fewer visually responsive and tuned neurons are excluded because they were not targeted for direct photostimulation.
e) The high-contrast trained decoders are then tested with low-contrast stimulus trials, yielding a range of success rates.
f) The performance of the decoder on low-contrast trials is related to the amount of correlation between the network activity patterns evoked by high and low contrast visual stimulation (Pearson correlation between the 1-x-n (n = number of ROIs) population vectors of concatenated single neurons average evoked response to the same orientation of either high or low stimulus contrast).
g) The decoder performs similarly regardless of whether the animal licked (no significant difference in decoder accuracy on hit compared to miss trials, $P = 0.068$ Wilcoxon signed rank paired, two-sided test).
h) The change in decoder accuracy on trials with photostimulation compared to trials without photostimulation (Wilcoxon signed rank paired, two-sided test with Bonferroni correction).
i) Relationship between the change in decoder accuracy and P(Lick) on the low contrast visual stimulus trials.
j) Relationship between the change in decoder accuracy and the associated change in P(Lick) on trials with compared to trials without photostimulation.
k) The average change in decoder accuracy when binarizing the change in P(Lick) into positive and negative groups (Wilcoxon signed rank paired, two-sided test).

**Supplementary Figure 9. Photostimulation evoked responses of target and follower cells on single trials in one example session.**

a) Stimulus triggered average traces of photostimulation responses, grouped by response type (directly targeted, excited, non-responsive, or inhibited). The thick line indicates the median across all cells and trials and the shaded region indicates the inter-quartile range. Dashed horizontal lines indicate the threshold used for classing a response in a given cell on a given trial as either excited or inhibited. The dotted vertical lines indicate the onset and offset of photostimulation.
b) Individual traces of all cells across all trials (sorted by response magnitude) grouped by whether the cell was directly targeted or if the response was classed as excited, non-responsive, or inhibited.
c) Illustration of the P(response) metric. Extracted responses are thresholded and each cell is then assigned a probability of response for both excitation and inhibition.
d) Comparison of the trial averaged P(response) metric for each cell in one example session with other cell average metrics (ΔF/F and AUC (which is related to the probability of a linear classifier correctly binarizing the distribution of baseline values from the distribution of response values)). The P(response) metric is highly correlated with both the ΔF/F metric and the AUC metric. Each point represents the average value across all trials for one cell. Dark coloured points indicate cells that are determined statistically, significantly responsive when comparing the distribution

of evoked responses to the distribution of baseline values with a Wilcoxon rank sum test (alpha=0.05).

**Supplementary Figure 10. Visual stimulus responsivity**

a) Stimulus triggered average activity traces during presentation of the cells preferred orientation of stimulus, in one example session. Cells can be either excited by (red, inhibited by (blue) or non-responsive (grey) to the set of presented stimuli. Thick lines indicate average (median) across all cells and shaded region indicated the inter quartile range across all cells.
b) Showing all individual cells, from one session, response to their preferred stimulus (that which elicits the maximum response), sorted by response magnitude and grouped by response type.
c) Tuning curves for each individual cell, in one example session, sorted by the direction of stimulus which produced the maximum response.
d) Average proportion of responsive cells, across all experiments.

**Supplementary Figure 11. Photostimulation ensembles within a session are parameter matched for spatial dispersion, size and responsivity but differ maximally in their responses to visual stimuli.**

a) The number of neurons within the 10 µm search radius around each SLM target location, that were detected as responsive to photostimulation.
b) The proportion of cells that were responsive to photostimulation in each session out of the number that were targeted.
c) The average photostimulation evoked response for each cell within each ensemble.
d) The spatial spread of targets within each ensemble is matched across ensembles. Measured by average pairwise distance between each target (d) and the average nearest neighbour distance (e).
e) The proportion of cells with each of the ensemble classes that were characterised as visually responsive.
f) The proportion of cell within each ensemble that share tuning to the fixed low-contrast stimulus orientation (selected to match the cotuned groups preference)
g) The average signal correlation, computed as the correlation coefficient between average tuning curves for each pair of cells, for each ensemble. All comparisons between groups made with Wilcoxon signed rank paired, two-sided test with Bonferroni correction. * = $P < 0.05/3$, ** = $P < 0.01/3$, *** = $P < 0.001/3$.

**Supplementary Figure 12. Example of the neuropil subtraction procedure.**

a) Example neuropil, ROI and neuropil-subtracted ROI traces from one cell. The segment of activity includes part of the orientation preference mapping block and the beginning of the photostimulation during behaviour block. Example photostimulation artefacts are indicated.
b) The correlation between the neuropil and the raw ROI signal (each point represents the average of 7 frames). The slope of the robust-regression fit line is used to scale the neuropil signal before subtracting it from the ROI signal. Dashed circles indicate both the set of points roughly corresponding to photostimulation artefacts (shared between the ROI and the neuropil) and ROI events (unique to the ROI, not seen in the neuropil).
c) The correlation between the neuropil and the neuropil-subtracted ROI signal.
d) The correlation between the raw ROI and the neuropil-subtracted ROI signal.

**Supplementary Figure 13. Minimal effects of neuropil subtraction on the spatial profile of the network response to photostimulation.**

a) The trial-averaged P(response) for excitation (left) inhibition (middle) and the difference between the two (right) for all cells, including directly targeted cells, binned spatially and aligned to the nearest target cell (0 µm lateral, 0 µm axial). The dashed vertical lines indicate the volume (20 µm diameter cylinder extending through all axial planes) excluded for 'follower' analysis in the manuscript. P(response) metrics calculated on raw, non-subtracted, ROI traces. Data at negative lateral displacements are the same as for positive displacements, for display purposes only.
b) Same as a) but for the final neuropil subtracted traces.

c) The average curves of P(response) against distance from nearest target for the lateral dimension at the 0 μm axial plane, and for the axial dimension at the 0 μm lateral position. Left: excitatory response, middle: inhibitory responses, right: the difference between the two. P(response) is calculated on the raw, non-subtracted, ROI traces.
d) Same as for c) but for the final neuropil subtracted traces.
e) Comparison of the lateral P(response) curves seen in c) and d) with and without neuropil subtraction. Shaded region indicates the standard deviation across all stimulation sessions. Left: excitatory response, middle: inhibitory responses, right: the difference between the two. ($N$ = 63 sessions, 14 mice. ** denotes $P < 0.01$, Wilcoxon signed rank test with Bonferroni correction).

**Supplementary Figure 14. Minimal effects of neuropil subtraction on the measured photostimulation response magnitude and detection of negative responses.**

a) Extracted photostimulation-triggered traces for all cells in one example experiment on one example trial. Solid line indicates the median across the cells in that response category (targeted, excited, non-responsive, inhibited). Shaded region indicates interquartile range. Dotted vertical lines indicate the photostimulation period, note this is the period affected by the photostimulation artefact and is the region with obvious difference between raw and neuropil subtracted traces. Responses are extracted in a window after photostimulation offset to avoid the artefact contamination.
b) Pooling all responses from all cells on all trials (to CT stimulation alone) in all sessions, comparing with and without neuropil subtraction.
c) Only considering the detected negative responses in the final, neuropil subtracted traces. All have a response of less than -1 $\Delta F/\sigma F$, as per criteria (shaded histogram). 95.4% of these negative responses are associated with a negative response even when not performing neuropil subtraction (white histogram).

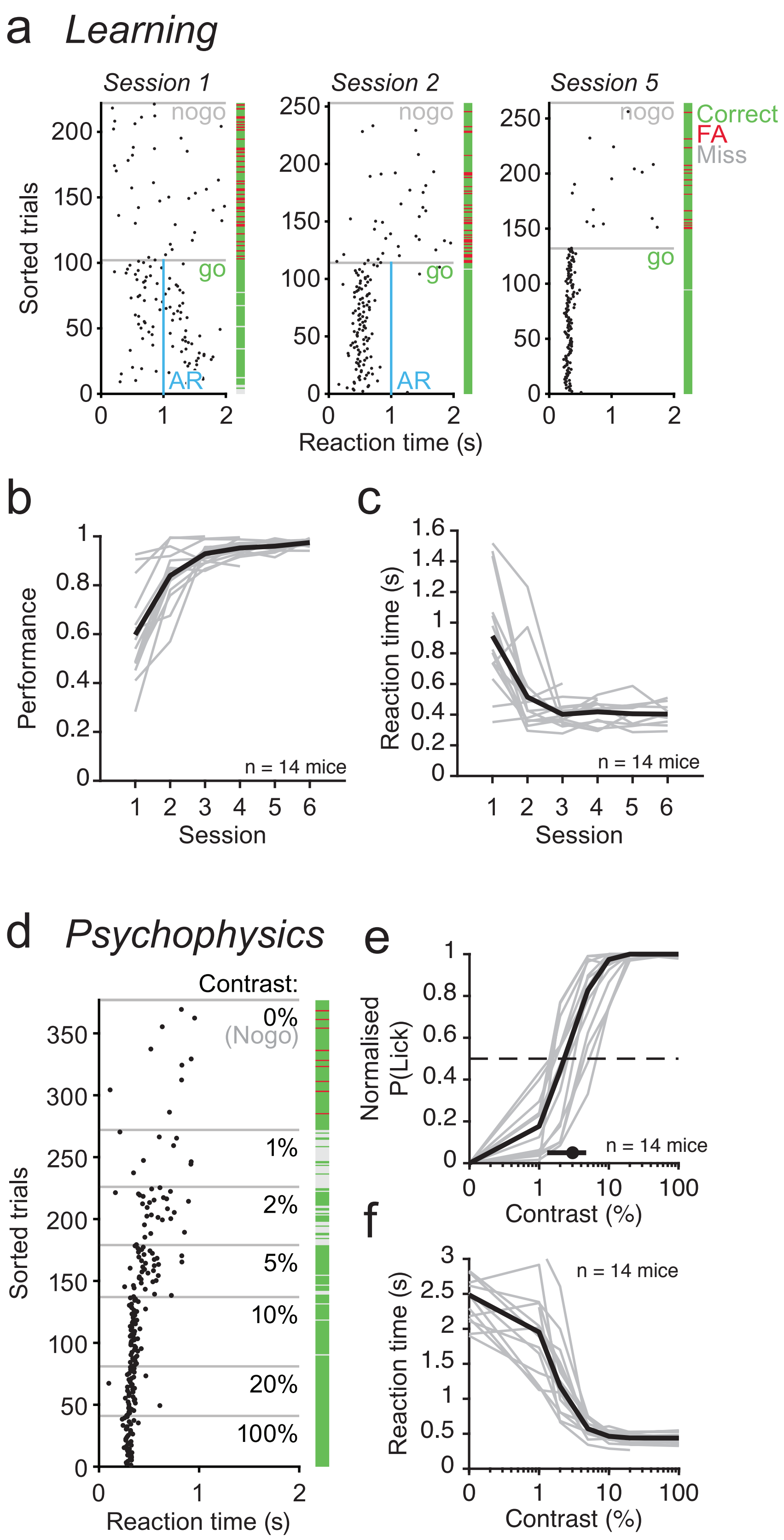


**Supplementary Figure 1.** Russell et al.

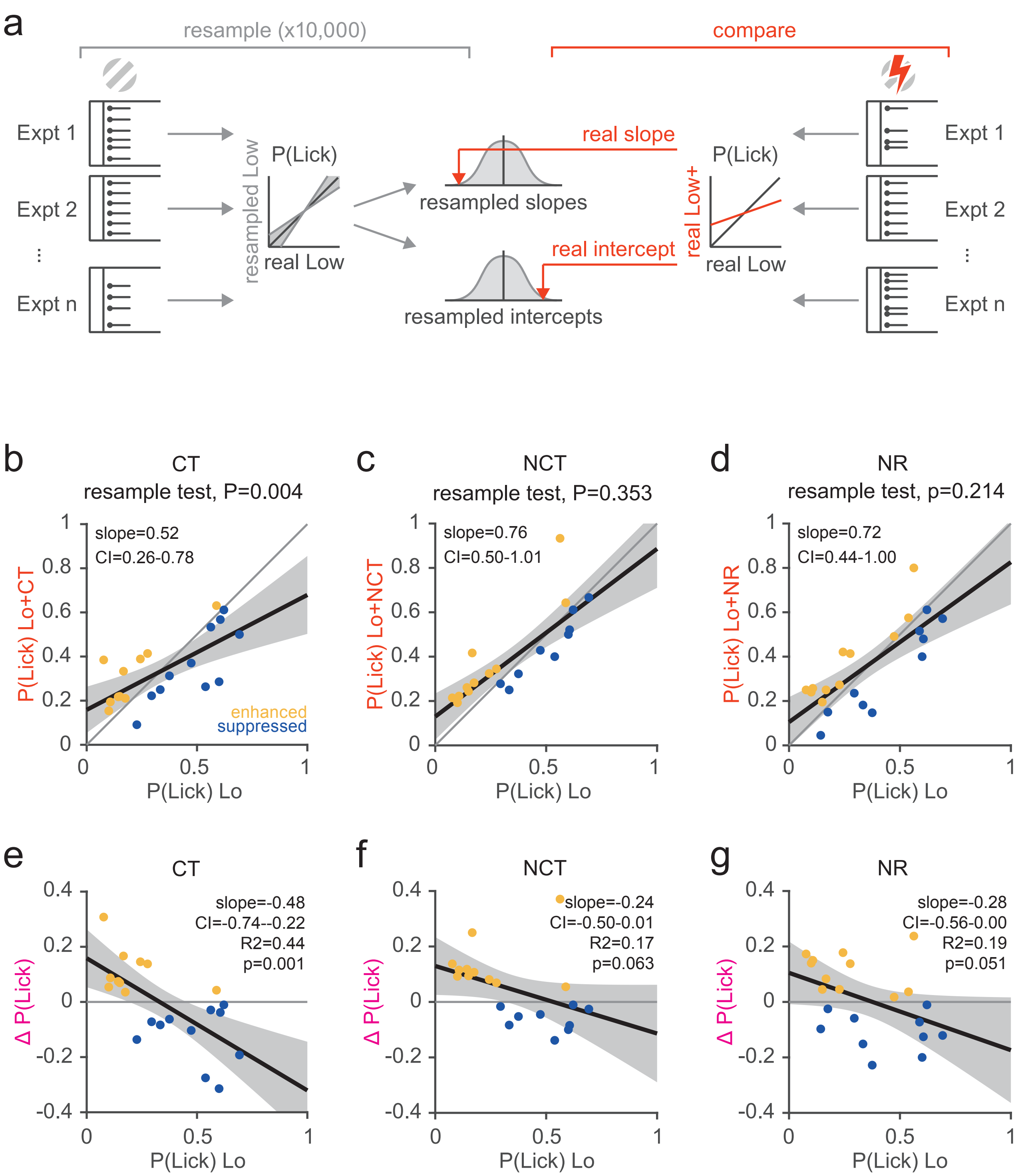


**Supplementary Figure 2.** Russell et al.

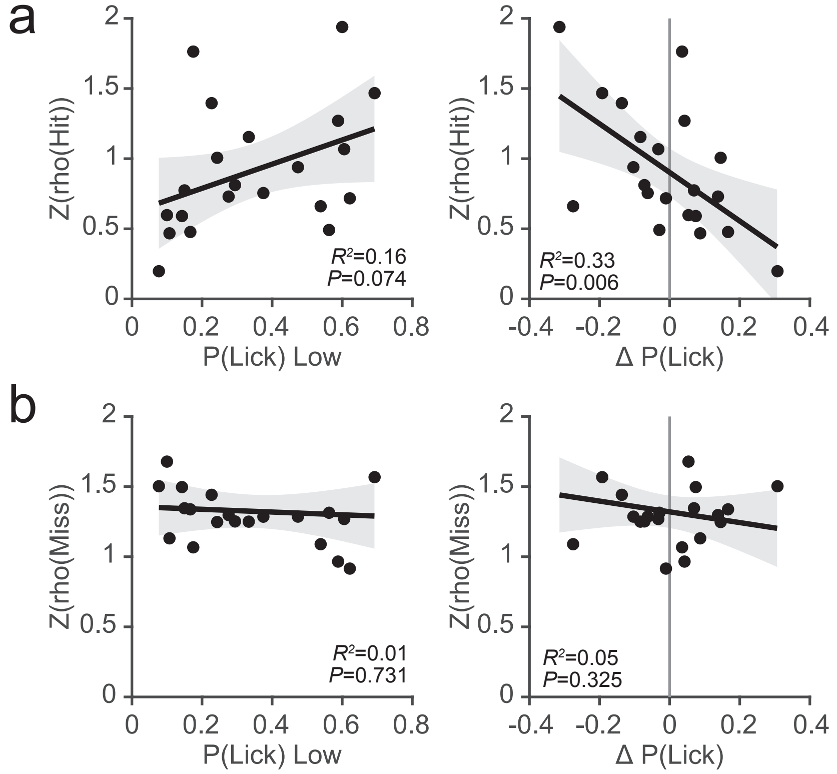


**Supplementary Figure 3.** Russell et al.

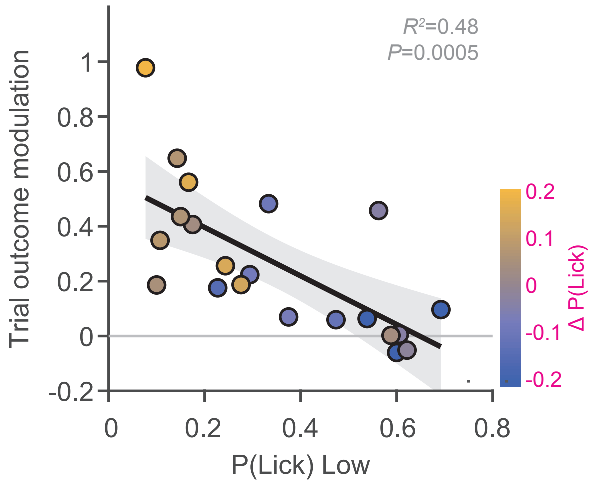


**Supplementary Figure 4.** Russell et al.

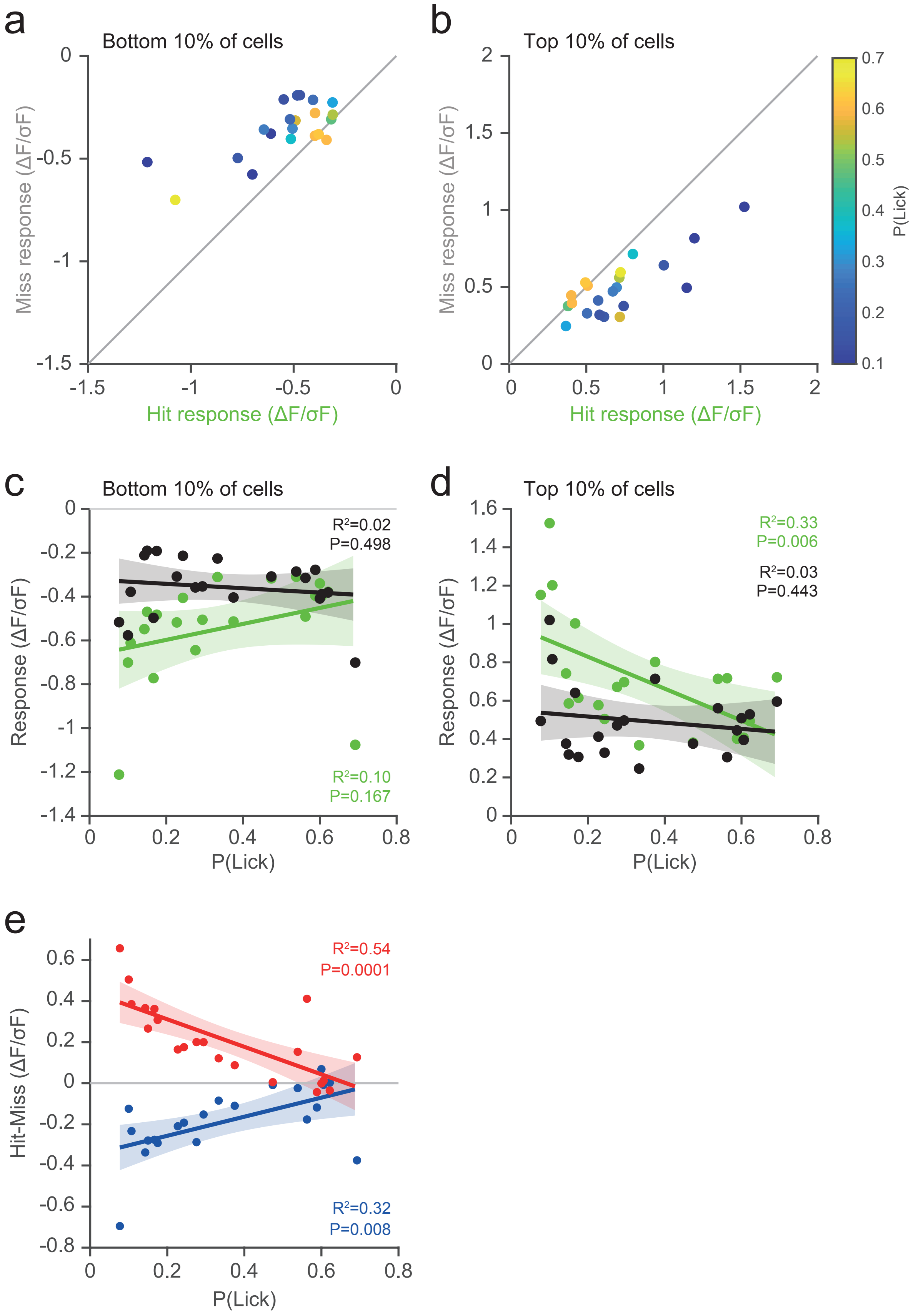


**Supplementary Figure 5.** Russell et al.

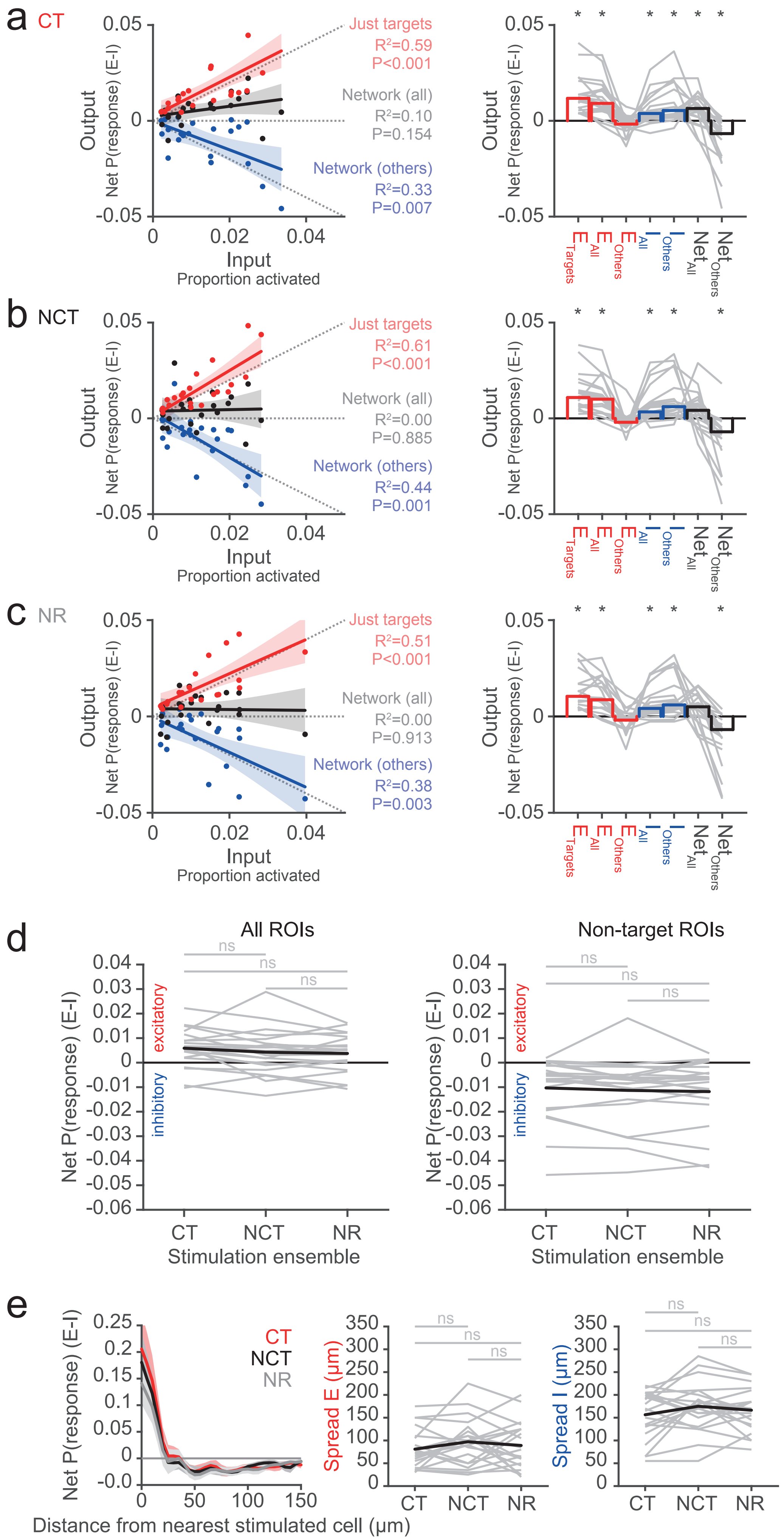


**Supplementary Figure 6.** Russell et al.

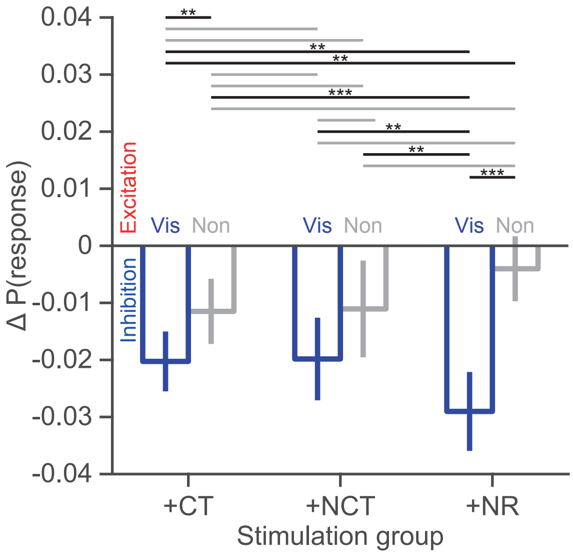


**Supplementary Figure 7.** Russell et al.

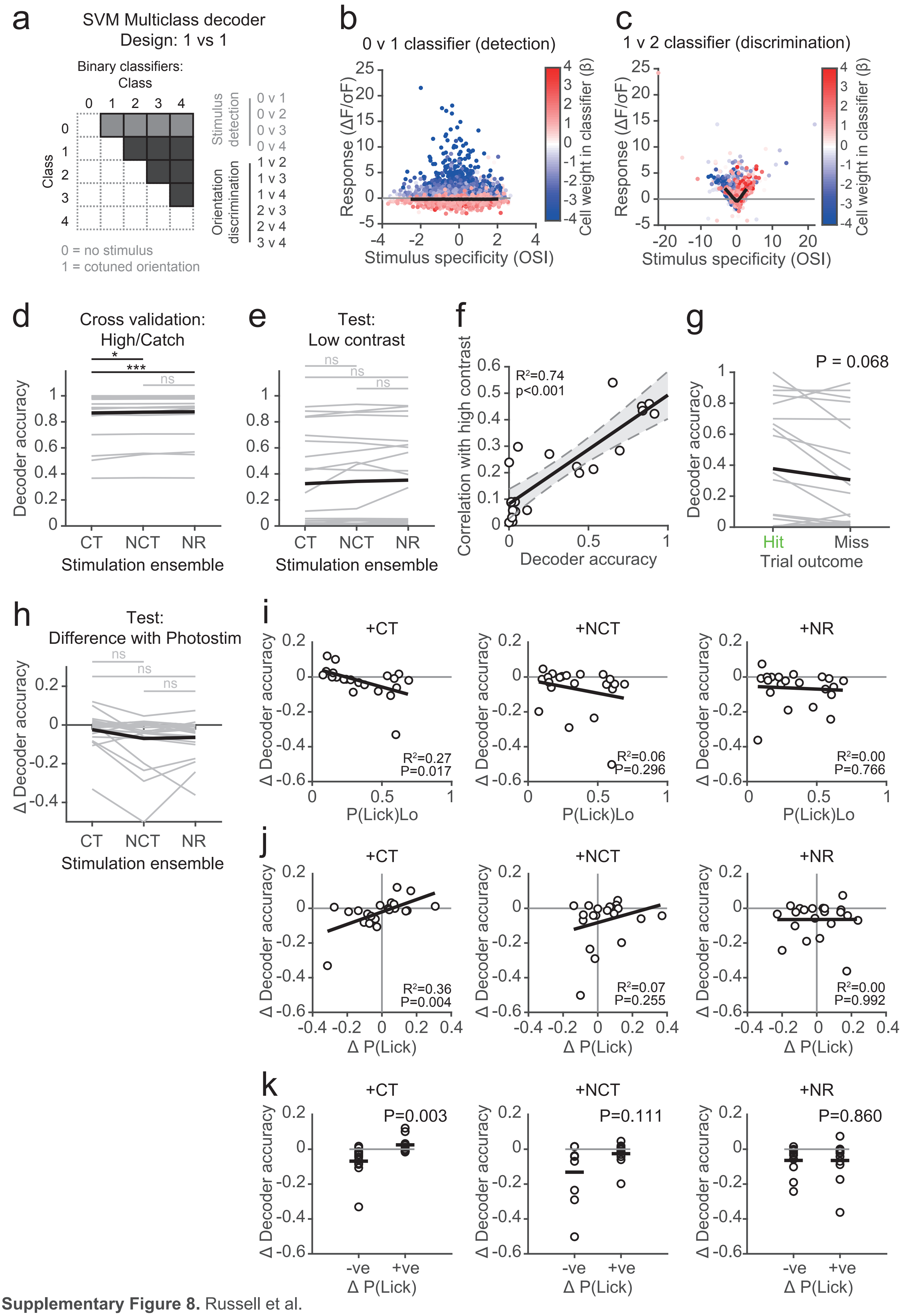


**Supplementary Figure 8.** Russell et al.

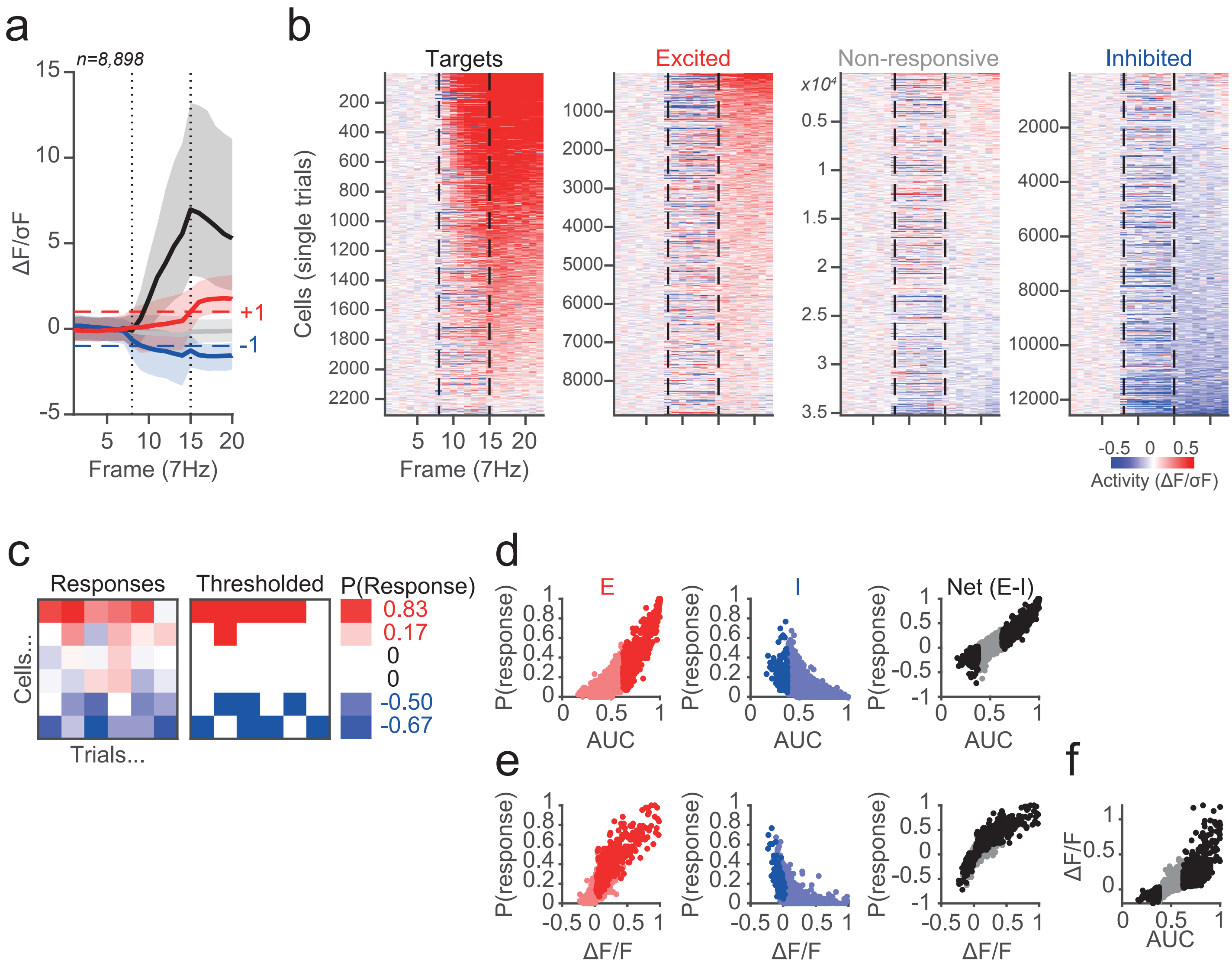


**Supplementary Figure 9.** Russell et al.

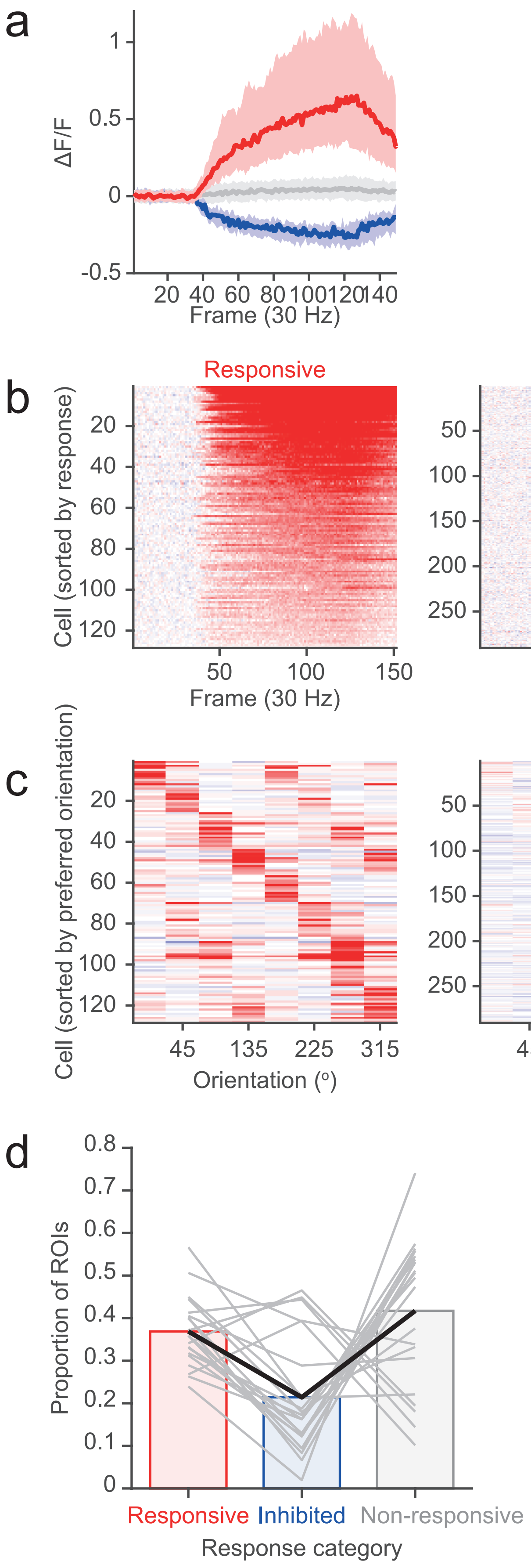


**Supplementary Figure 10.** Russell et al.

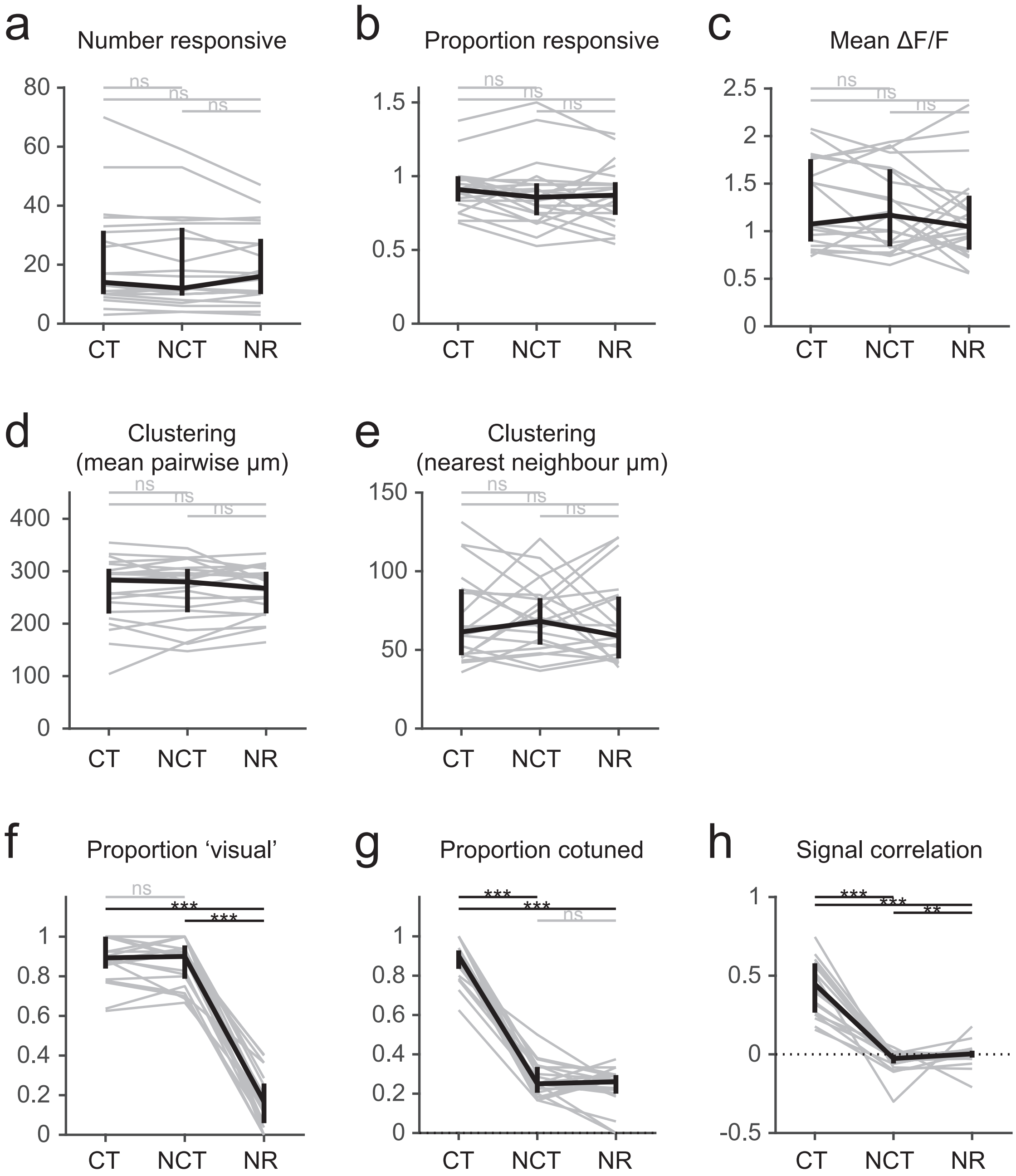


**Supplementary Figure 11.** Russell et al.

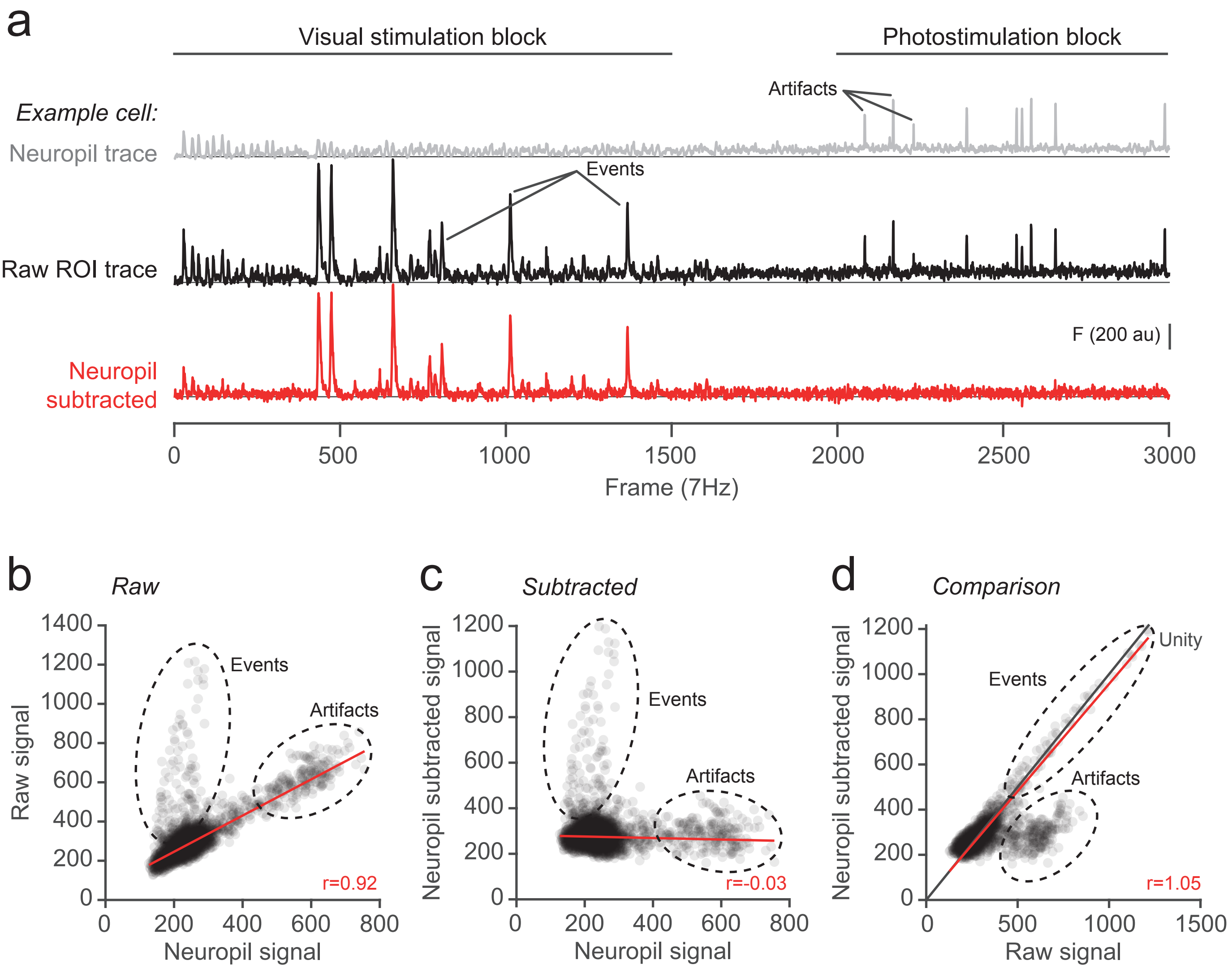


**Supplementary Figure 12.** Russell et al.

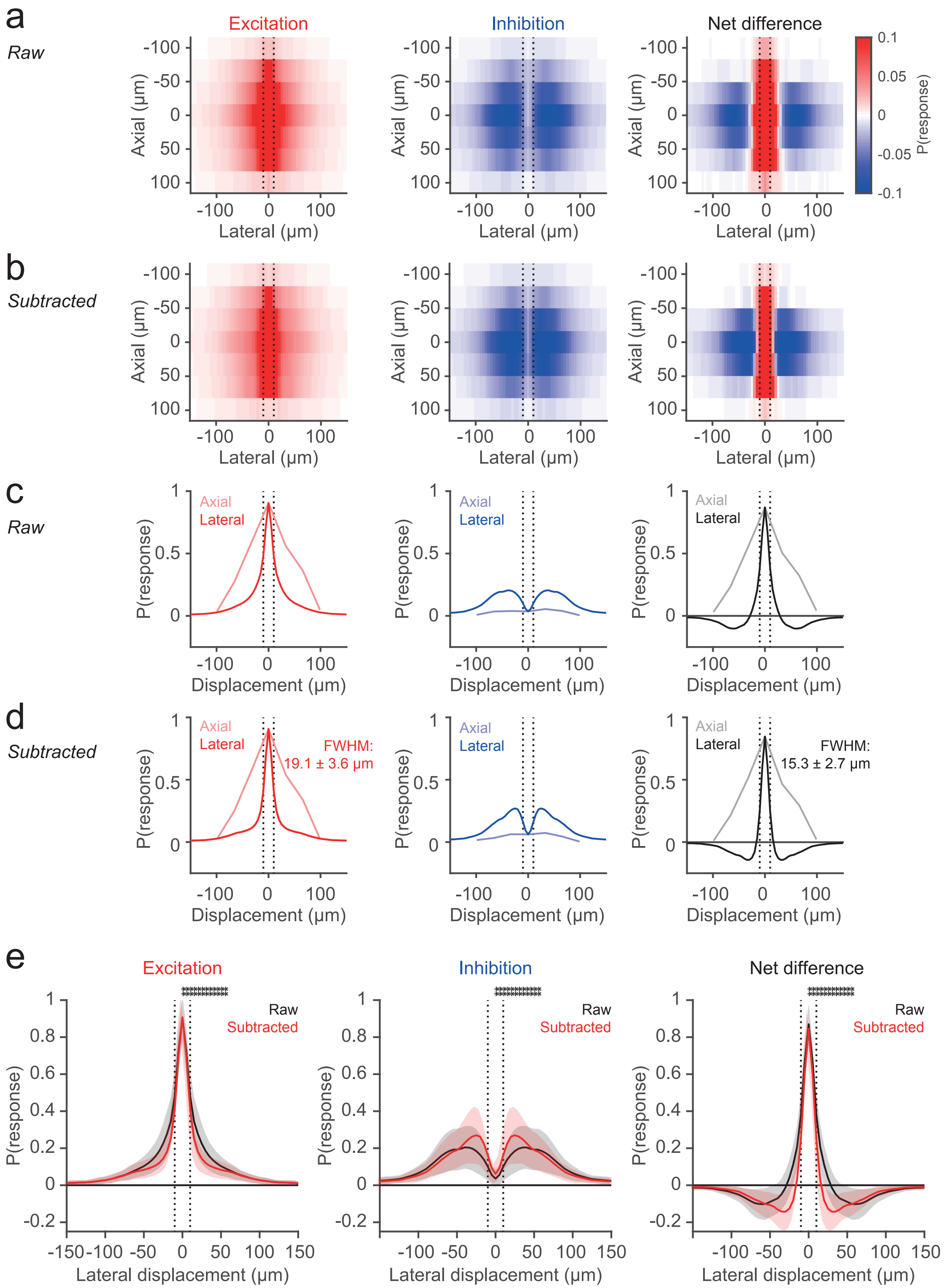


**Supplementary Figure 13.** Russell et al.

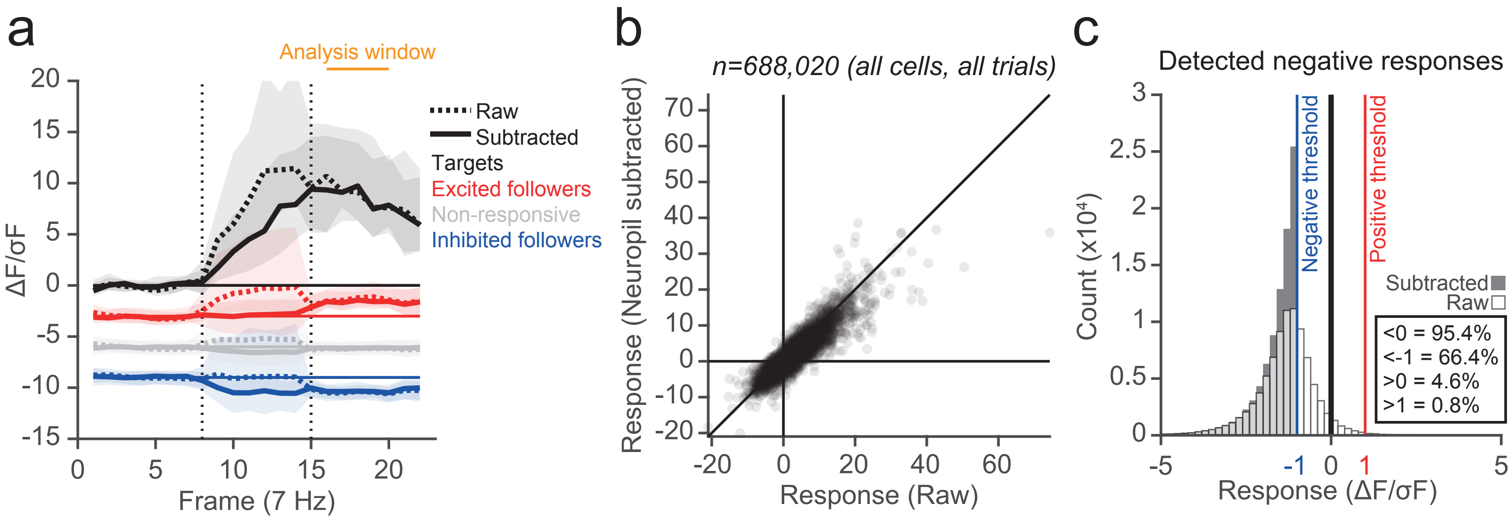


**Supplementary Figure 14.** Russell et al.
